# A small signaling domain controls PPIP5K phosphatase activity in phosphate homeostasis

**DOI:** 10.1101/2024.09.12.612650

**Authors:** Pierre Raia, Kitaik Lee, Simon M. Bartsch, Felix Rico-Resendiz, Daniela Portugal-Calisto, Oscar Vadas, Vikram Govind Panse, Dorothea Fiedler, Michael Hothorn

**Author notes:** Center for Structural Biology, Center for Cancer Research, National Cancer Institute (NCI), Frederick, MD 21702-1201, U.S.A.

## Abstract

Inositol pyrophosphates (PP-InsPs) are highly phosphorylated nutrient messengers. The final step of their biosynthesis is catalyzed by diphosphoinositol pentakisphosphate kinase (PPIP5K) enzymes, which are conserved among fungi, plants, and animals. PPIP5Ks contain an N-terminal kinase domain that generates the active messenger 1,5-InsP_8_ and a C-terminal phosphatase domain that participates in PP-InsP catabolism. The balance between kinase and phosphatase activities controls the cellular levels and signaling capacity of 1,5-InsP_8_. Here, we present crystal structures of the apo and substrate-bound Vip1 phosphatase domain from S. *cerevisiae* (ScVip1^PD^). ScVip1^PD^ is a phytase-like inositol 1-pyrophosphate phosphatase with two conserved histidine phosphatase catalytic motifs. The enzyme has a strong preference for 1,5-InsP_8_ and is inhibited by inorganic phosphate. ScVip1^PD^ has an α-helical insertion domain stabilized by a structural Zn^2+^ binding site, and a unique GAF signaling domain that exists in an open and closed state, allowing channeling of the 1,5-InsP_8_ substrate to the active site. Mutations that alter the active site, that restrict the movement of the GAF domain or that modify the charge of the substrate channel, significantly inhibit the activity of the yeast enzyme *in vitro*, and the function of the Arabidopsis PPIP5K VIH2 *in planta*. Structural analyses of full-length PPIP5Ks suggest that the kinase and phosphatase are independent enzymatic modules. Taken together, our work reveals the structure, enzymatic mechanism and regulation of eukaryotic PPIP5K phosphatases.

## Introduction

Inositol pyrophosphates (PP-InsPs) are eukaryotic nutrient messengers^1^. These intracellular signaling molecules consist of a densely phosphorylated six-carbon *myo*-inositol ring with pyrophosphate groups attached, for example, at position 1 (1-InsP_7_), position 5 (5-InsP_7_), or both positions (1,5-InsP_8_). PP-InsPs regulate cell signaling by a variety of mechanisms, including mediating protein-protein interactions, allosterically regulating enzyme or transporter activities, or post-translationally modifying proteins^2,3^.

A central role of PP-InsPs in cellular inorganic phosphate (Pi) homeostasis is conserved among fungi^4–6^, protozoa^7^, algae^8^, plants^9–15^, and animals^16–19^. Pi homeostasis is primarily controlled by 1,5-InsP_8_ binding to PP-InsP sensing SPX receptor proteins^4,5,11,12,14,15,17^. 1,5-InsP_8_ levels are low under Pi starvation conditions, whereas 1,5-InsP_8_ accumulates under Pi-sufficient growth conditions or upon Pi resupply^5,13^. As cellular Pi homeostasis and other essential cellular processes appear to be mechanistically linked to 1,5-InsP _8_ levels, understanding the regulation of 1,5-InsP_8_ biosynthesis and catabolism is of fundamental importance.

PP-InsPs are generated from InsP_6_ through a series of phosphorylation steps catalyzed by different small molecule kinases. In yeast, the inositol hexakisphosphate kinase Kcs1^20,21^ and in plants, the inositol 1,3,4-trisphosphate 5/6-kinases ITPK1/2^22^, preferentially transfer a second phosphate to position 5 of InsP_6_ or 1-InsP_7_ to generate 5-InsP_7_ and 1,5-InsP_8_, respectively. The diphosphoinositol pentakisphosphate kinases (PPIP5Ks), known as Vip1 in *S. cerevisiae*, Asp1 in *Schizosaccharomyces pombe*^23–25^, PPIP5K in amoeba and in animals^16,26–30^ or VIP/VIH in plants^8,10,31–34^, preferentially transfer a phosphate to position 1, to generate 1-InsP_7_ and 1,5-InsP_8_, respectively. Consistent with the function of 1,5-InsP_8_ as messenger in Pi homeostasis in Arabidopsis, *vih1/vih2* loss-of-function mutants have severely reduced 1,5-InsP_8_ levels and display constitutive Pi starvation responses^10–12^.

In addition to the conserved Pi homeostasis-related phenotypes, loss-of-function of PPIP5K enzymes results in altered growth, cell and vacuolar morphology, chromosome segregation, host colonization and DNA damage-induced autophagy^24,25,35,36^. PPIP5K mutants are associated with hearing loss in humans and mice ^37^. In human cell lines, depletion of PPIP5K activity induced broad changes in cell metabolism^38^. In plants, PPIP5K activity has been linked to jasmonic acid and salicylic acid hormone signaling, and to cell wall architecture^32–34,39^.

There are several PP-InsP catabolic enzymes, including a family of histidine acid phosphatases located in C-terminal half of PPIP5Ks^24^. Thus, PPIP5Ks are dual-function enzymes that contain two distinct domains with opposing enzymatic activities: An N-terminal kinase domain that preferentially generates 1,5-InsP_8_ from 5-InsP_7_^25,26,40^, and a C-terminal phosphatase domain with inositol 1-pyrophosphate phosphatase activity^25,29,40–42^. The relative PP-InsP kinase and phosphatase activities in PPIP5K are thought to modulate the cellular levels of 1,5-InsP_8_ ^2,10,16^.

PPIP5K loss-of-function mutants show altered PP-InsP pools in various eukaryotes^5,8,10,13,24,25,32,34–36,38,43^. PPIP5K knockout mutants have reduced 1-InsP_7_ and 1,5-InsP_8_ levels, while the 5-InsP_7_ substrate of the PPIP5K kinase domain accumulates^5,13,34,44^. 1,5-InsP_8_ pools can be restored to wild-type like levels in genetic rescue experiments with phosphatase-dead but not with kinase-dead versions of the corresponding PPIP5K enzyme^10,25,35,36,38^.

How regulation of PPIP5Ks controls 1,5-InsP_8_ pools in response to changes in Pi nutrient availability is not well understood. Currently, it is known that the ATP/ADP ratio decreases in response to Pi starvation, which in turn reduces 1,5-InsP_8_ synthesis by the kinase domains of human PPIP5K1/2 or plant VIH1/2^10,16^. Pi itself has been reported as an inhibitor of the phosphatase domain of human and fungal PPIP5Ks^10,16,40^.

The structure and enzymatic mechanism of the human and fungal PPIP5K kinase domains have been well established^27,45,46^. However, currently there is no structural information on the PPIP5K phosphatase domain and its enzymatic mechanism and regulation remain to be elucidated. Based on sequence comparison, the PPIP5K phosphatase domain has been assigned to the histidine acid phosphatase superfamily^24^, specifically to the branch 2, which includes the well-characterized *Escherichia coli* and fungal phytases, as well as glucose-1-phosphate, and acid phosphatases^47^. The PPIP5K phosphatase domain contains the catalytic RHxxR motif, which is conserved among histidine acid phosphatases^24^. However, the lack of sequence conservation in the substrate-binding domains within the superfamily precludes predictions about substrate recognition and specificity. A “cryptic” pleckstrin homology (PH) domain located near the conserved RHxxR motif has been hypothesized to play a critical role in substrate binding ^26^. PH domains, typically found in signaling proteins, bind phospholipids and molecules derived from phospholipid headgroups, including inositol phosphate and PP-InsPs^48^. We now report the structure of a PPIP5K phosphatase domain and characterize its substrate binding and enzymatic mechanism, its regulation by nutrients and its structural organization within the full-length enzyme.

## Results

### Overall structure of the ScVip1 kinase and phosphatase domains

We first attempted to characterize the structure and catalytic mechanism of the histidine acid phosphatase domain (PD) from plant PPIP5Ks^10,31,32^. However, we were unable to express sufficient amounts of the isolated phosphatase domains from either *Arabidopsis thaliana* VIH1/VIH2 or from *Marchantia polymorpha* VIP1^34^ (Supplementary Fig. 1a). Therefore, we expressed and purified the phosphatase domains from *S. cerevisiae* Vip1 (ScVip1^PD^, residues 536-1107) as well as its N-terminal kinase domain (ScVip1^KD^, residues 186-522) in baculovirus-infected insect cells (see Methods).

The structure of the ScVip1^KD^ was solved by molecular replacement to a resolution of ∼1.2 Å (Table 1). Apo crystals of the ScVip1^PD^ initially diffracted to ∼3.1 Å. The structure was solved by molecular replacement - single wavelength anomalous diffraction (MR-SAD) on a platinum derivative (Table 1). In this structure, a large disordered loop (residues 851-918) was located and subsequently replaced by a short Gly-Ser-Ser-Gly linker. Crystallization of ScVip1^PDΔ848-918^ yielded a second crystal form diffracting at 3.4 Å resolution (Table 1).

**Table 1.**
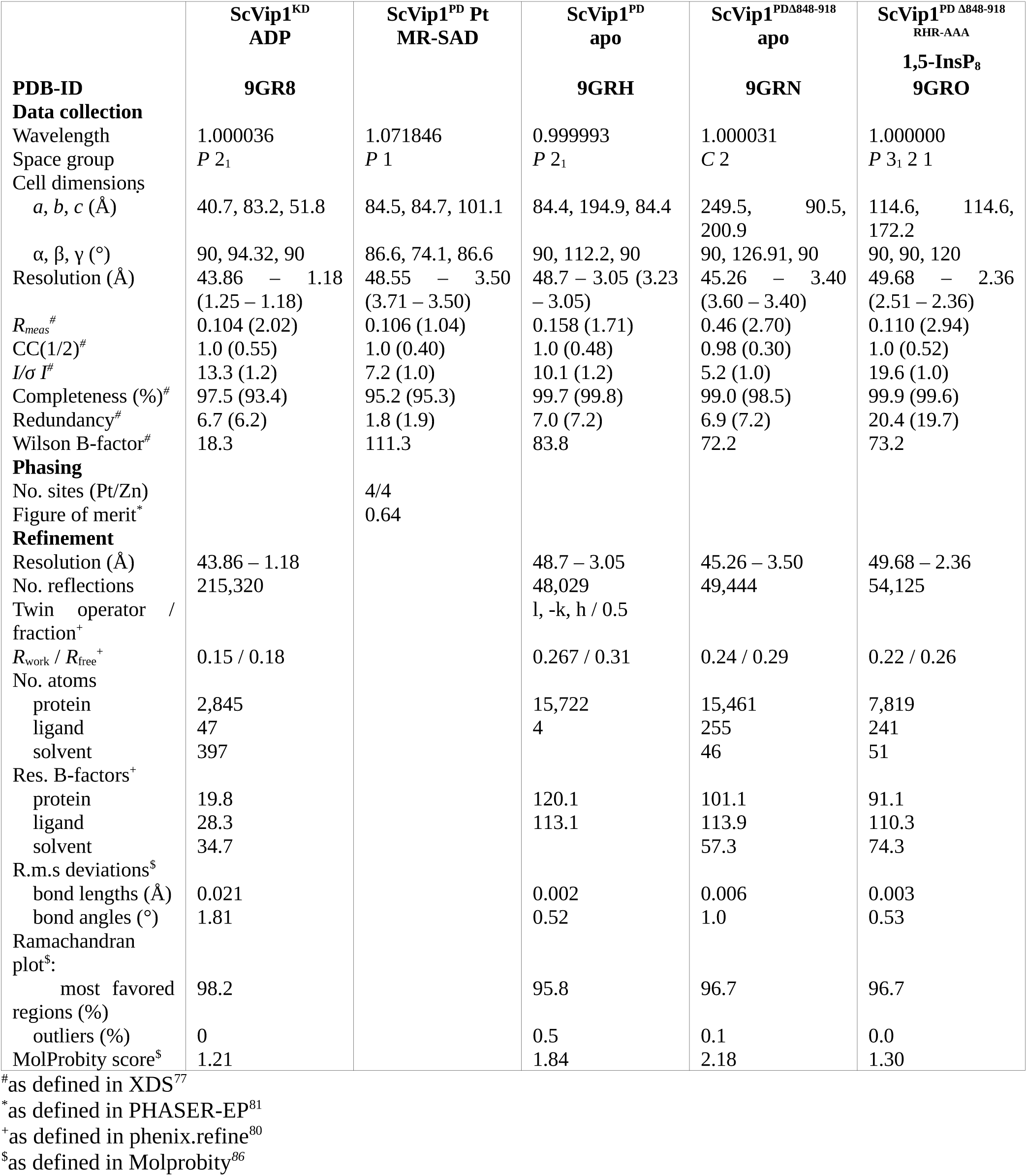
Crystallographic data collection and refinement statistics.

The structure of ScVip1^KD^ in complex with adenosine diphosphate revealed the typical ATP-grasp fold and substrate binding sites previously seen in other PPIP5K structures^27,46^ (Fig. 1a, b). ScVip1^KD^ shares significant sequence and structural homology with HsPPIP5K2 and with the kinase domain of *S. pombe* Asp1 (Supplementary Figs. 2, 3). The C-terminal α11 helix of the ScVip1 kinase domain (residues 498-520) is well defined in our structure (Fig. 1b; Supplementary Fig. 2).

**Fig. 1.**
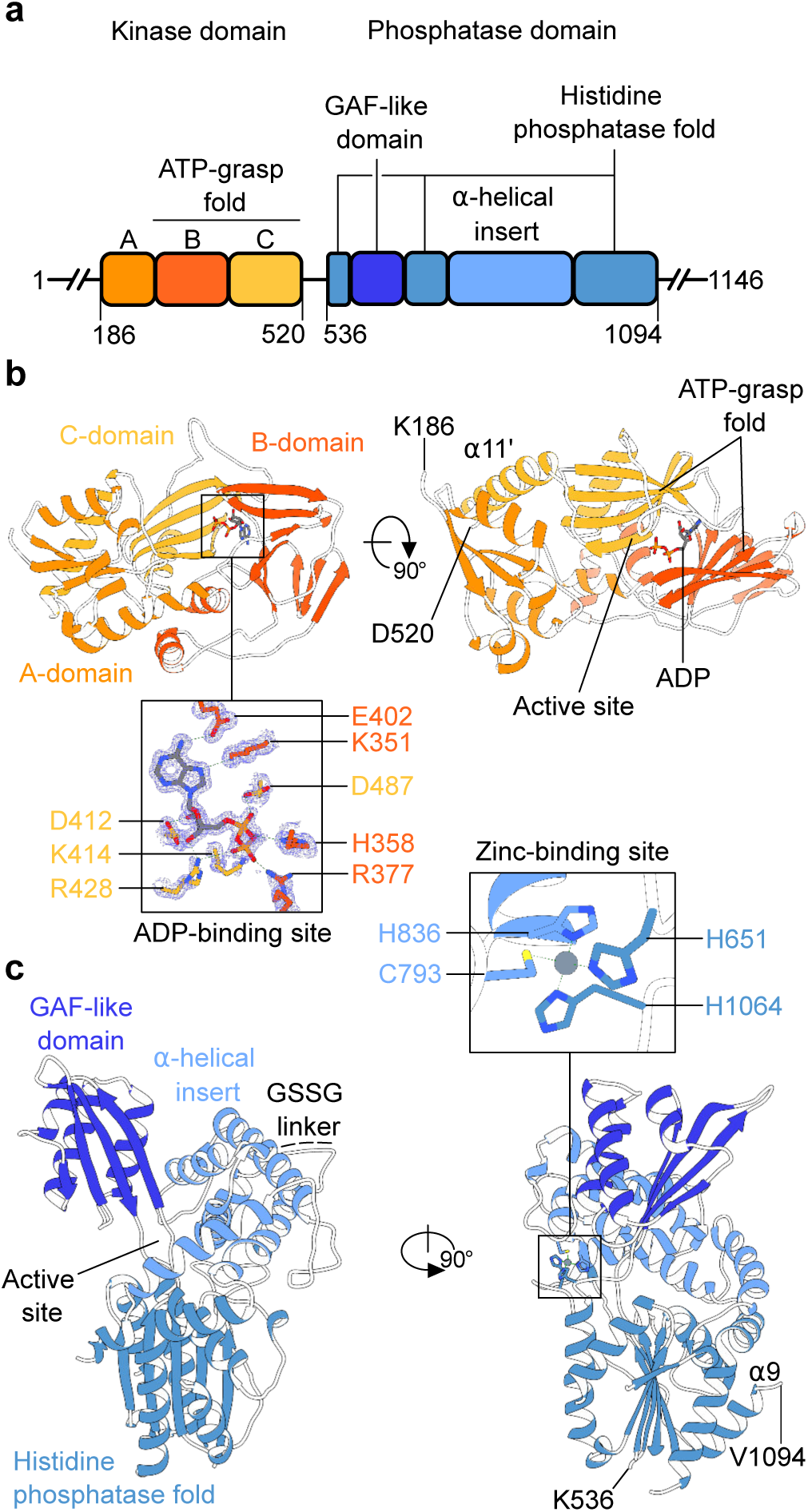
PPIP5K enzymes contain an unusual histidine acid phosphate domain. **a** Schematic diagram of the *S. cerevisiae* Vip1 kinase (orange, ScVip1^KD^) and phosphatase (blue, ScVip1^PD^) domains (https://www.uniprot.org/ Uniprot ID Q06685), highlighting structured regions as blocks (residues 186-520 and 536-1094) and putative linker regions as solid lines (residues 1-185, 521-535, and 1095-1146). **b** Ribbon diagram of the ScVip1^KD^ structure bound to ADP (in bonds representation) and including a omit 2 (*F_o_ – F_c_*) electron density map contoured at 1 σ (blue mesh). **c** Ribbon diagram of ScVip1^PD Δ848-918^, with the histidine acid phosphatase core and α-helical insertion domain shown in light blue, and the PPIP5K-specific GAF domain in dark blue, respectively. The position of the engineered GSSG linker is indicated with a dotted line. The inlet provides a view of the zinc binding site, with the Zn ^2+^ ion shown as a sphere (in gray) and the coordinating histidine and cysteine residues depicted in bonds representation.

The two apo structures of ScVip1^PD^ revealed a conserved histidine acid phosphatase core with two insertion domains (Fig. 1c). The N-terminus of ScVip1^PD^ (residue 536) and the most C-terminal α9 helix (residues 1079-1094) are well defined by electron density (Fig. 1c). Based on these domain boundaries, we hypothesize that the ScVip1 kinase and phosphatase domains are surrounded by large, unstructured N- and C-terminal extensions, and are connected by a flexible ∼15 amino acid linker (Fig. 1a; Supplementary Fig. 2, see below). Plant and human PPIP5Ks contain sequence-diverse linkers of similar length between their kinase and phosphatase domains (Supplementary Fig. 2). An unexpected metal binding site occupied by a Zn^2+^ ion is formed by His651 and His1064 from the histidine acid phosphatase core, and by Cys793 and His836 from the α-helical insertion domain (Fig. 1c; Supplementary Fig. 2; see below).

### A PPIP5K unique GAF domain

Next, we performed structural homology searches with ScVip1^PD^ as a search model using the DALI web server (http://ekhidna2.biocenter.helsinki.fi/dali/)^49^. We found that ScVip1^PD^ shares the core phosphatase domain fold and the α-helical insertion domain with bacterial, plant and human histidine acid phosphatases (Fig. 2a; Supplementary Fig. 4). The relative orientation of the α-helical insertion domain is conserved among histidine acid phosphatases with very different substrate preferences (Fig. 2a), including phytases that hydrolyze InsP_6_ (Fig. 2b). We located a second insertion domain specific to PPIP5Ks in a position normally occupied by the much smaller β1-insert in other histidine acid phosphatases (Fig. 2a). Analysis with DALI revealed a weak structural homology of this second insertion domain with bacterial and human GAF ^50^ or PAS^51^ domains involved in light, metabolite and cyclic nucleotide sensing^52,53^ (Fig. 2c; Supplementary Fig. 4). The GAF domain is conserved among fungal, plant and animal PPIP5Ks (Supplementary Fig. 2). The central β-sheet of the ScVip1^PD^ GAF domain is reminiscent of the inositol phosphate / inositol polyphosphate binding surface previously identified in PH domains, but its topology is different (Fig. 2c) ^54,55^.

**Fig. 2.**
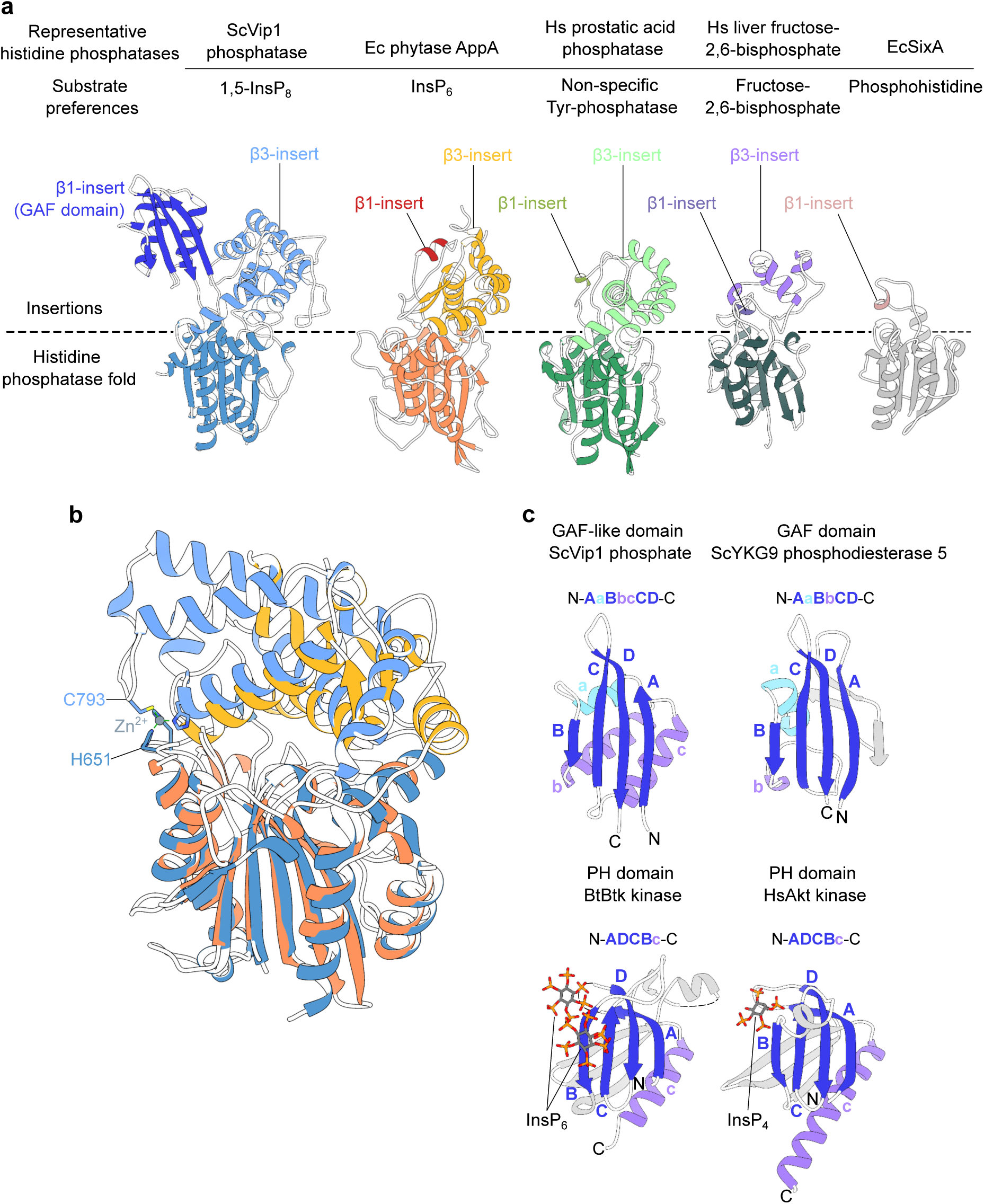
A unique GAF domain in PPIP5Ks replaces the canonical β1-insert in histidine acid phosphatases. **a** Structural comparison of ScVip1^PD Δ848-918^ (ribbon diagram, in blue) with the *E. coli* phytase AppA (https://www.rcsb.org/ PDB-ID 7Z2S^60^, yellow to red), human prostatic acid phosphatase (PDB-ID 1ND6^99^, light- to dark-green), human liver fructose-2,6-bisphosphatase (PDB-ID 1K6M^71^, purple and dark gray) and the sensor histidine kinase phosphatase SixA from *E. coli* (PDB-ID^100^ 1UJC, pink and light gray). **b** Structural superposition of ScVip1^PD Δ848-918^ and EcAppA (colors as in panel A, root mean square deviation r.m.s.d. is ∼ 1.1 Å comparing 105 corresponding C_α_ atoms from the histidine acid phosphatase core). The Zn^2+^ binding site in ScVip1^PD^ is not conserved among other histidine acid phosphatases. **c** Structural comparison of the GAF domain in ScVip1^PD^ and structurally related GAF (PDB-ID 1F5M^52^, r.m.s.d. is ∼0.9 Å comparing 26 corresponding C_α_ atoms in the β-sheet) and inositol polyphosphate-binding PH domains (PDB-IDs 4Y94^55^, 1UNQ^54^). Note that GAF and PH domains share similar sized β-sheets with different topologies.

### ScVip1^PD^ specifically hydrolyses 1,5-InsP_8_

1,5-InsP_8_ is the preferred substrate for the isolated ScVip1 phosphatase domain in quantitative nuclear magnetic resonance (NMR) phosphatase assays using ^13^C-labeled substrates (Fig. 3a; Supplementary Fig. 5). ScVip1^PD^ is a specific inositol pyrophosphate phosphatase, with a high preference for the 1-position of PP-InsPs (Fig. 3a; Supplementary Fig. 5), as previously reported^25^. In malachite green-based phosphatase assays, we derived similar *K*_M_ values for 1,5-InsP_8_ and 1-InsP_7_, but the catalytic efficiency of ScVip1^PD^ for 1,5-InsP_8_ is ∼18fold higher when compared to 1-InsP_7_ (Fig. 3b). Notably, 1,5-InsP_8_ is the reaction product of the N-terminal kinase domain of ScVip1^24^.

**Fig. 3.**
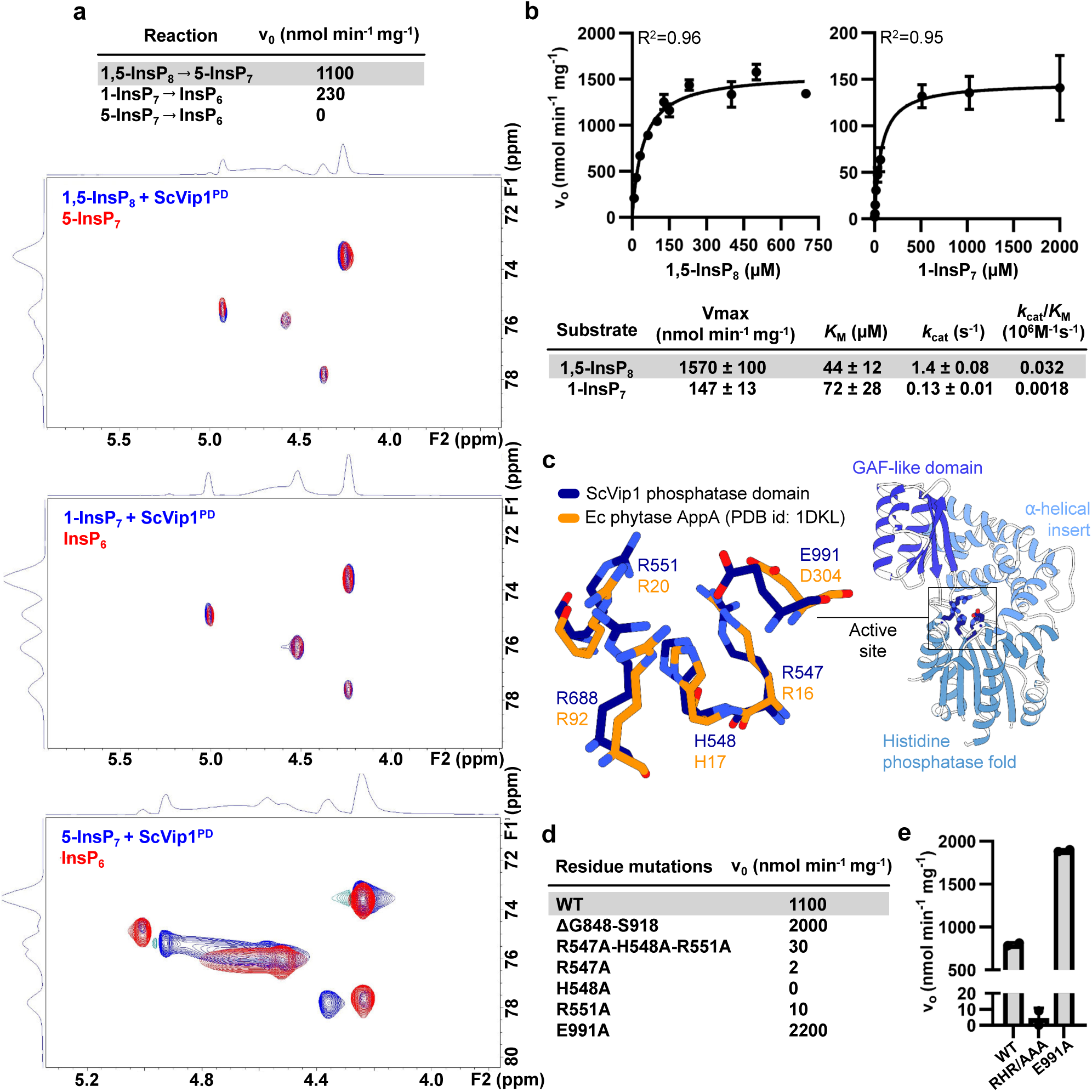
ScVip1^PD^ specifically hydrolyzes 1,5-InsP_8_ using a conserved phytase active site. **a** Table summary of the initial enzyme velocities (v_0_) for *Sc*Vip1^PD^ versus 1,5-InsP_8_, 1-InsP_7_, or 5-InsP7 derived from an NMR-based phosphatase assay. Pseudo-2D NMR spectra of ScVip1^PD^ incubated with different substrates (blue traces) are shown below, overlaid with the respective reaction products 5-InsP_7_, or InsP_6_ (red traces). **b** Initial enzyme velocities (v_0_) for ScVip1^PD^ – catalyzed hydrolysis of 1,5-InsP_8_ and 5-InsP_7_ as a function of inositol pyrophosphate concentration and using a malachite green-based phosphatase assay. The Michaelis-Menten equation was fitted to the data using non-linear regression. Extracted kinetic parameters are summarized in the table as means ± S.E.M. (n=2). **c** Structural superposition of the active sites of ScVip1^PD Δ848-918^ (in bond representation, in dark blue) and EcAppA (in orange, PDP-ID 1DKL, r.m.s.d. is ∼1.0 Å comparing 67 corresponding C_α_ atoms from the histidine acid phosphatase core). In EcAppA, His17 is the catalytic residue, Asp304 is the proton-donor, and Arg16, Arg20, and Arg92 are involved in InsP_6_ substrate binding. **d-e** Initial velocities of 1,5-InsP_8_ hydrolysis catalyzed by wild-type and mutant versions of ScVip1^PD^, measured by NMR-based (**d**) or malachite green-based (**e**) phosphatase assays.

A sequence motif central to catalysis in histidine acid phosphatases is located in the active site cleft of ScVip1^PD^ sandwiched between the phosphatase core and the α-helical insertion domain (Fig. 3c). The motif with the consensus sequence Arg-His-x-x-Arg (RHxxR, where x can be any amino acid) is structurally conserved between ScVip1^PD^ and bacterial phytases (Fig. 3c). Mutation of either Arg547, His548 or Arg551 to alanine greatly reduces 1,5-InsP_8_ hydrolysis in NMR- and malachite green-based enzyme assays, respectively (Fig. 3d, e; Supplementary Fig. 5). This is consistent with the established function of the two arginine residues in InsP_6_ substrate binding in bacterial phytases^56^, and the essential role of the nucleophilic acceptor histidine in catalysis^57^. Asp304 in *E. coli* phytase is the proton donor involved in the degradation of the phosphohistidine intermediate^58^. In our ScVip1^PD^ structures, a glutamate residue (Glu991) occupies the position of the phytase proton donor aspartate (Fig. 3c). Mutation of Glu911 to alanine activates the enzyme (Fig. 3d, e; Supplementary Fig. 5, see below). Similarly, deletion of the large disordered loop in the α-helical insertion domain (ScVip1^PDΔ848-918^) moderately activates the engineered enzyme *in vitro* (Fig. 3d).

In conclusion, the isolated phosphatase domain of ScVip1 is an inositol 1-pyrophosphate phosphatase with a substrate preference for 1,5-InsP_8_ and with an active site similar to InsP_6_ metabolizing phytases.

### Inhibition of ScVip1^PD^ by Pi

Gu et al. previously reported that low millimolar concentrations of Pi inhibited the inositol 1-pyrophosphate phosphatase activity of human PPIP5K1 and PPIP5K2^16^. Addition of 10 mM Pi inhibited the phosphatase activity of ScVip1^10^. We found that ScVip1^PD^ is inhibited by Pi in a concentration-dependent manner, as inferred from NMR-based assays (Fig. 4a). As previously reported for SpAsp1 ^40^, sulfate was less effective than Pi in inhibiting the ScVip1^PD^ phosphatase activity (Fig. 4b). Nitrate had no detectable effect on the hydrolysis of 1,5-InsP_8_ (Fig. 4c). We conclude that Pi at concentrations found in the yeast cytosol ^59^, directly inhibits the enzymatic activity of the isolated ScVip1 phosphatase domain.

**Fig. 4.**
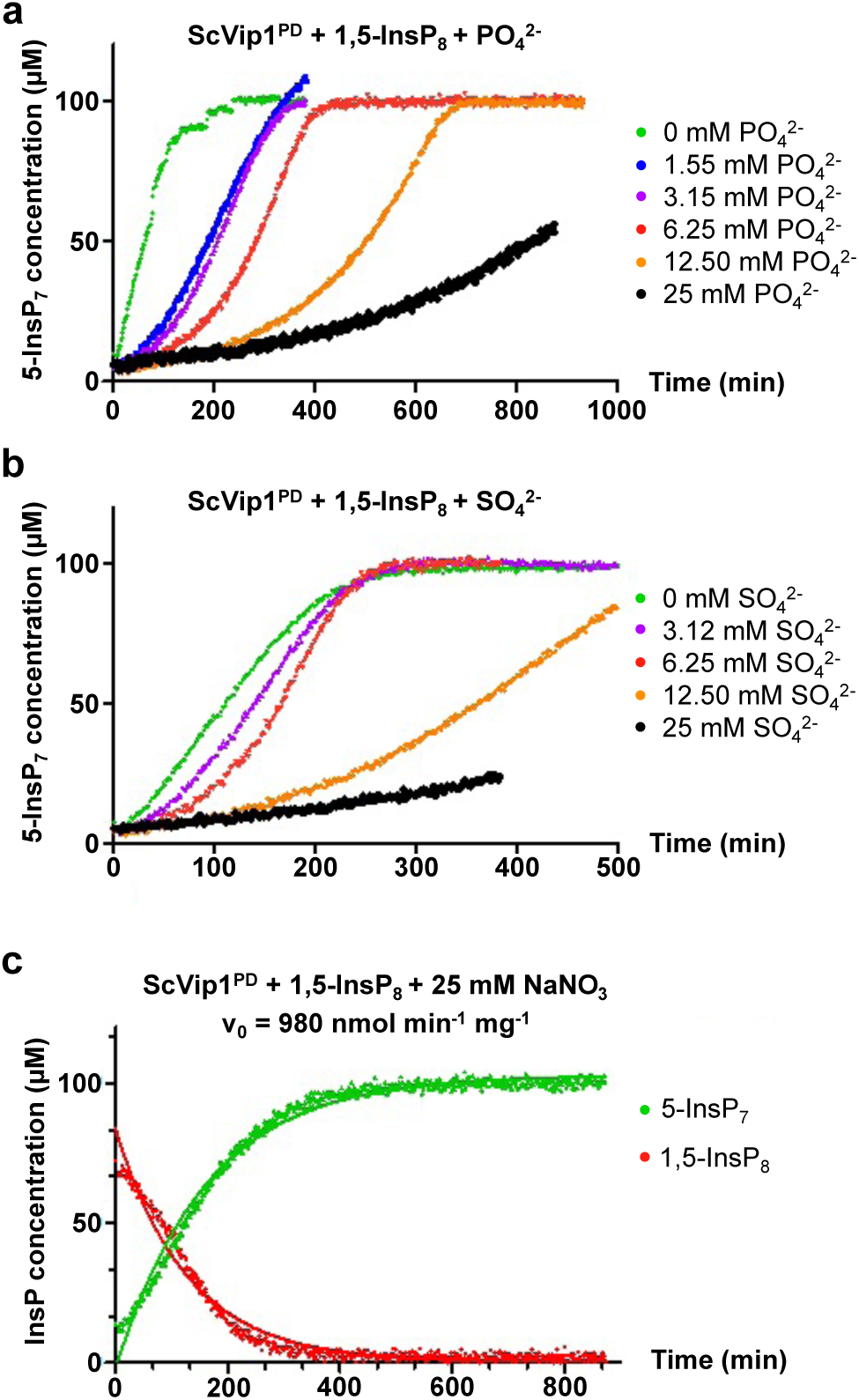
ScVip1^PD^ phosphatase activity is inhibited by inorganic phosphate and sulfate. **a-b** NMR time course experiments for ScVip1^PD^-catalyzed hydrolysis of 100 µM 1,5-InsP_8_ to 5-InsP_7_ in the presence of increasing concentrations of (**a**) PO ^2-^, (**b**) SO ^2-^, or (**c**) at a constant concentration of 25 mM NaNO_3_. The measured initial velocity of 980 nmol min^-1^ mg^-1^ is similar to the initial velocity in absence of NaNO_3_ (1100 nmol min^-1^ mg^-1^, compare Fig. 3a).

### GAF domain movements control enzyme activity

All four molecules in the asymmetric unit of our ScVip1^PD^ crystals display the GAF domain in an “open” conformation (Fig. 5a; Supplementary Fig. 6; Table 1). In ScVip1^PDΔ848-918^ crystals two molecules in the asymmetric unit also adopt the “open” conformation (Fig. 5a; Supplementary Fig. 6; Table 1). Crystal packing interactions between two adjacent GAF domains in our ScVip1^PDΔ848-918^ crystals are mediated in part by a second Zn^2+^ binding site. The Zn^2+^ ion is coordinated along a pseudo two-fold axis involving residues His573 and Glu575 from the GAF domain (Supplementary Fig. 6). Given that ScVip1^PD^ behaves as a monomer in solution (Supplementary Fig. 7) and that the histidine and glutamate residues are not conserved among other PPIP5Ks (Supplementary Fig. 2), the second Zn^2+^ coordination site is likely a crystallization artifact (Supplementary Fig. 6).

**Fig. 5.**
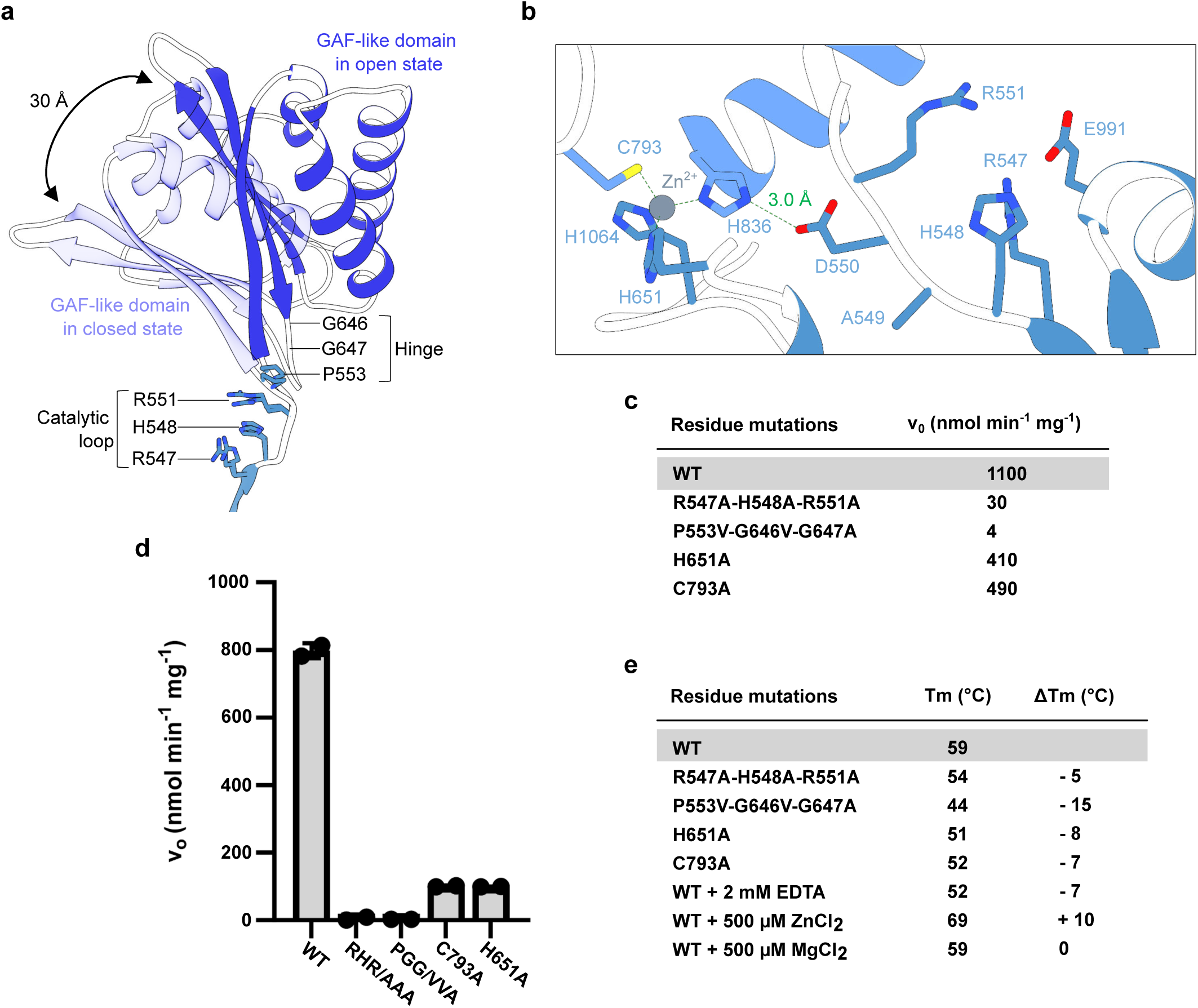
ScVip1^PD^ is regulated by GAF domain movements and stabilized by a zinc-binding site. **a** Structural superposition of apo ScVip1^PD Δ848-918^ reveals the GAF insertion domain in either its open (chain A, dark blue ribbon diagram) or closed (chain D, in light blue) conformation. The RHxxR and hinge regions are shown alongside (in bonds representation) **b** Details of the ScVip1^PD Δ848-918^ active site: Asp550 stabilize the catalytic loop by engaging in a salt bridge (green dotted line) with the Zn ^2+^ ion coordinating His836. The active site and Zn^2+^ coordinating residues are depicted in bonds representation, the Zn^2+^ ion is shown as a gray sphere. **c-d** Initial velocities of 1,5-InsP_8_ hydrolysis catalyzed by wild-type or mutant version of ScVip1^PD^, measured by NMR (**c**) or malachite green (**d**) phosphatase assays. **e** Table summarizing the thermal melting profiles of wild-type or mutant ScVip1^PD^. The ΔTm (in °C) represent the difference between the melting temperature of a designated ScVip1^PD^ mutant protein and the wild-type control.

Next, we replaced the catalytic loop residues Arg547, His548 and Arg551 in ScVip1^PD Δ848-918^ with alanine and obtained crystals in complex with the substrate 1,5-InsP_8_ diffracting at ∼2.4 Å resolution (ScVip1^PD Δ848-918 RHR-AAA^, Table 1). Notably, the GAF domain adopts a “closed” conformation in the substrate-bound structure (Fig. 5a). We hypothesized that the movements of the GAF domain are enabled by a hinge region in close proximity to the catalytic loop (RHxxR motif) of the enzyme (Fig. 5a, b). Consequently, mutation of the hinge residues Pro553, Gly646 and Gly647 (P553V-G646V-G647A) inactivated ScVip1^PD^ in NMR- and malachite green-based phosphatase assays, respectively (Fig. 5b-d). This suggests that movement of the GAF domain is part of the enzymatic cycle of ScVip1^PD^. Notably, the catalytic loop (Arg547, His548, Arg551) does not appear to be involved in the domain movement when comparing the open and closed conformation structures (Fig. 5a). The conserved Zn^2+^ binding site in ScVip1^PD^ positions His836 to form a salt bridge with Asp550, apparently stabilizing the position of the catalytic loop (Fig. 5b). Mutation of the Zn^2+^ binding site residues His651 or Cys793 to alanine reduces the phosphatase activity in our enzyme assays, suggesting that the Zn^2+^ binding site is involved in the structural stabilization of the enzyme and, importantly, of its catalytic loop (Fig. 5b-e). Consistent with this, mutation of Zn ^2+^ coordinating residues reduces the structural stability of ScVip1^PD^, whereas excess of ZnCl_2_ further stabilizes the enzyme in thermal shift assays *in vitro* (Fig. 5e, Supplementary Fig. 7).

### The GAF domain is involved in substrate binding and channeling

In our ScVip1^PD Δ848-918 RHR-AAA^ – 1,5-InsP_8_ complex structure, we found a domain-swapped crystallographic dimer stabilized by an intermolecular disulfide bridge (Supplementary Fig. 8). One molecule of 1,5-InsP_8_ is bound to each monomer, either on the surface of the enzyme involving the GAF domain or near the catalytic center (Fig. 6a; Supplementary Fig. 8). The GAF domain adopts a closed conformation in both monomers (Fig. 6a). In both binding sites, all phosphate groups of 1,5-InsP _8_ are well defined by electron density (Fig. 6b), with the 1-β-phosphate facing away from the active site (Fig. 6c). Conserved basic residues from the GAF domain and the α-insertion domain contribute to the formation of the outer substrate binding surface (Fig. 6c). A large network of lysine residues from the GAF domain and from the base of the α-insertion domain forms the second substrate binding surface, which is located in close proximity to the RHxxR catalytic loop (Fig. 6c). Most of the basic residues contributing to the outer and the inner 1,5-InsP_8_ binding surfaces map to the central β-sheet of the GAF domain (Fig. 6c, d). Comparison of the outer and inner substrate binding sites in ScVip1^PD^ suggests that the 1,5-InsP_8_ substrate may undergo a ∼ 90° rotation upon entering the active site (Fig. 6d).

**Fig. 6.**
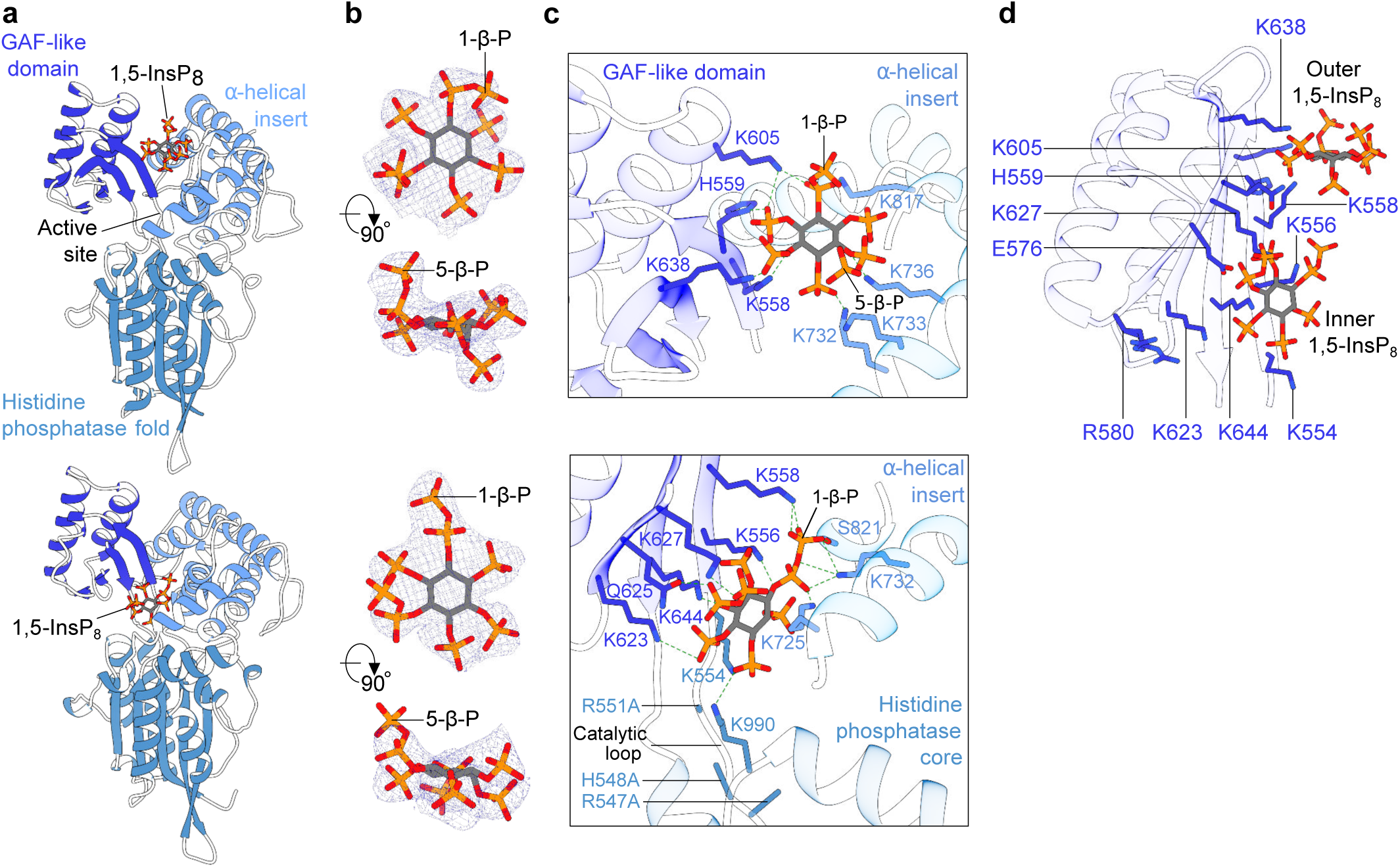
The GAF domain of ScVip1^PD^ binds 1,5-InsP_8_ in two capture sites. **a** Ribbon diagrams of ScVip1^PD Δ848-918 RHR-AAA^ bound to 1,5-InsP_8_ (in bonds representation). 1,5-InsP_8_ binds either at the surface of ScVip1^PD Δ848-918 RHR-AAA^ (chain A, top) or close to the active site (chain B, bottom). **b** Detailed view of the 1,5-InsP_8_ molecules shown in two orientations and including a 2 (*F_o_ – F_c_*) electron density map (blue mesh) contoured at 2.5 σ in chain A (top) or at 1 σ in chain B (bottom), respectively. “1-β-P” and “5-β-P” designate the positions of the β-phosphate of the pyrophosphate group attached to the position 1 or 5 of the *myo*-inositol ring. **c** Detailed view of the 1,5-InsP_8_ binding sites. The residues interacting with 1,5-InsP_8_ are shown in bonds representation, hydrogen bonds are indicted by dotted lines (in green). **d** Ribbon diagram of the isolated GAF domain from the ScVip1^PD Δ848-918 RHR-AAA^ structure, depicting the relative positions of the outer (chain A) and inner (chain B, modeled) 1,5-InsP_8_ substrate (in bonds representation) binding sites.

We next compared the open conformation of apo ScVip1^PD^ with the substrate-bound structures (Fig. 7a). We found that the conformational changes strongly alter the surface charge distribution of the enzyme, with the GAF domain opening and closing on a large and well conserved basic surface area surrounding the substrate binding sites and formed by the GAF and α-insertion domains (Fig. 7a,b). By mapping the positions of the 1,5-InsP_8_ molecules bound to the outer and inner substrate binding sites, we uncovered a potential substrate binding channel (Fig. 7c). We hypothesize that the 1,5-InsP_8_ substrate first targets the outer substrate binding site of ScVip1^PD^ in its open conformation. Substrate binding to this outer site may trigger closure of the GAF domain, resulting in rotation of the substrate and its translocation to the active site (Fig. 7c). In its closed conformation, the PPIP5K GAF domain contributes to the formation of a substrate channel that may facilitate transport of 1,5-InsP_8_ to the catalytic center of the enzyme as well as the exit of the 5-InsP_7_ product from the enzyme (Fig. 7c).

**Fig. 7.**
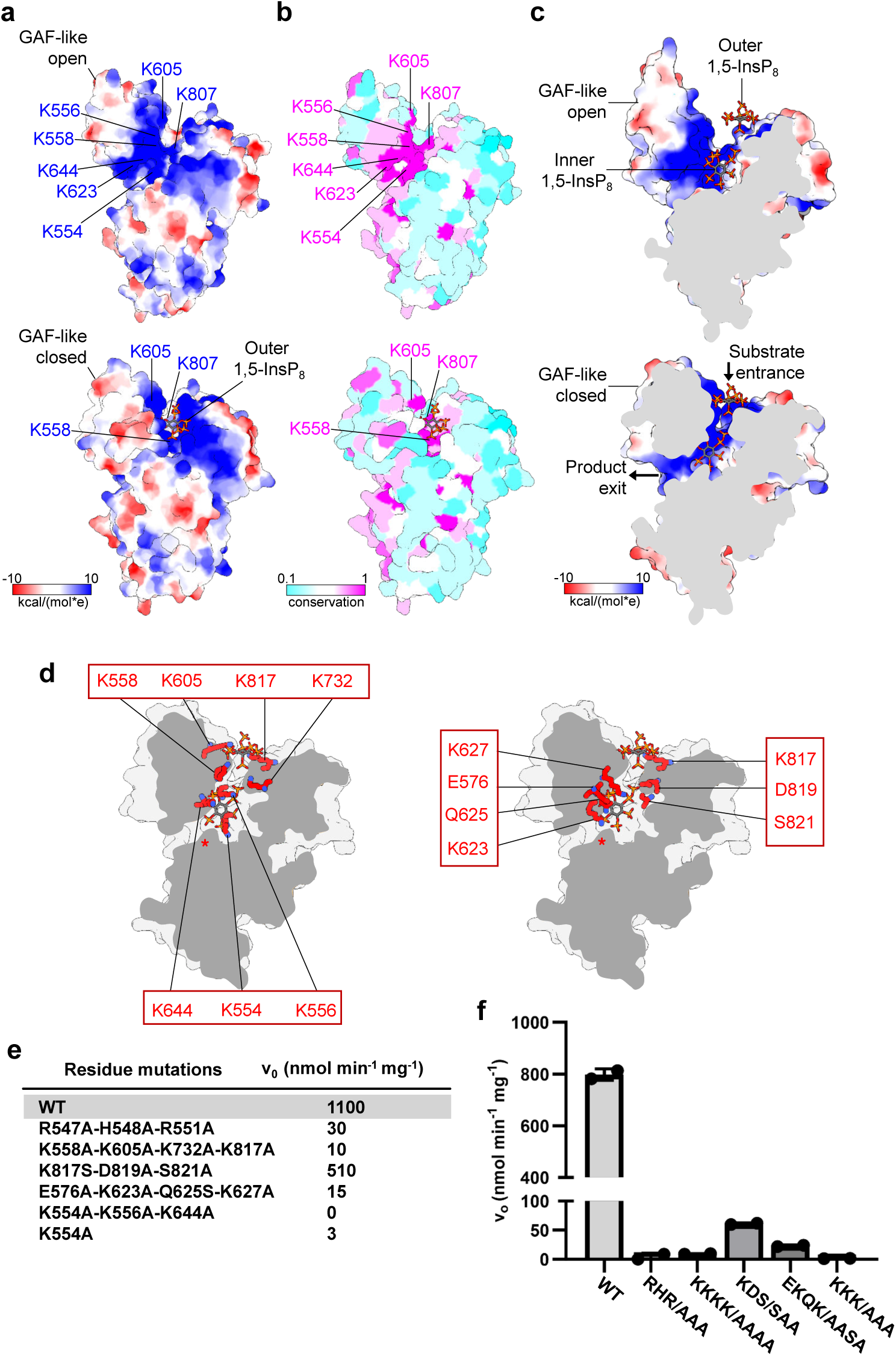
A GAF domain-induced substrate channel transports 1,5-InsP_8_ to the active site. **a** Electrostatic surface charge distribution of the apo ScVip1^PD Δ848-918^ (top) and 1,5-InsP_8_-bound ScVip1^PD Δ848-918 RHR-AAA^ (bottom) structures. **b** Evolutionary conservation (from cyan, variable to pink, invariant) of surface residues mapped onto the apo ScVip1^PD Δ848-918^ (top) and 1,5-InsP_8_-bound ScVip1^PD Δ848-918 RHR-AAA^ (bottom) structures. Views as in **a**. Residue conservation was derived from the multiple sequence alignment shown in Supplementary Fig. 2 and analyzed with Consurf^101^. **c** Cutaway front view of the apo ScVip1^PD Δ848-918^ surface modeled with the outer and inner 1,5-InsP_8_ molecules (top). Same view for the 1,5-InsP_8_-bound ScVip1^PD Δ848-918 RHR-AAA^ surface modeled with the outer and inner 1,5-InsP_8_ molecules (bottom). The closed GAF domain forms the substrate channel. **d** Groups of mutated residues involved in 1,5-InsP_8_ channeling (in bonds representation) mapped onto the ScVip1^PD Δ848-918 RHR-AAA^ structure (shown as molecular surface). The active site of ScVip1^PD^ is indicated with a red asterisk. **e-f** Initial velocities of 1,5-InsP_8_ hydrolysis for wild-type or mutant versions of ScVip1^PD^, measured by NMR-based (**e**) or malachite green-based (**f**) phosphatase assays.

Mutation of highly conserved residues from the outer substrate binding site (Lys558, Lys605, Lys732, Lys817 to alanine) (Fig. 7d) inactivated the enzyme (Fig. 7e, f), suggesting that 1,5-InsP_8_ is indeed initially bound to the surface of the enzyme before entering the active site. We next mutated residues that contribute to the PPIP5K substrate channel. Mutation of Lys817, Asp819 and Ser821 originating from the α-helical insertion domain (Fig. 7d) moderately reduced the 1,5-InsP_8_ phosphohydrolase activity of ScVip1^PD^ in both NMR- and malachite green-based assays (Fig. 7e, f). In contrast, mutation of 1,5-InsP_8_ binding residues from the GAF domain (Glu576, Lys623, Gln625, Lys627) inactivated the enzyme (Fig. 7d-f). Mutations close to the active site of the enzyme (Lys554, Lys557, Lys644) also reduced the activity of ScVip1 ^PD^ (Fig. 7d-f), but in addition structurally destabilized the enzyme (Supplementary Fig. 7). We therefore generated an additional single Lys554 to alanine mutant, which is structurally intact (Supplementary Fig. 7) but which also showed very low enzyme activity in our assays (Fig. 7d-f). Taken together, our structure-function analyses suggest that PPIP5K phosphatase domains contain two 1,5-InsP_8_ binding sites that contribute to substrate channeling into the active site.

### An active site glutamate mediates substrate specificity of ScVip1^PD^

As described above, the canonical phytase proton donor aspartate is replaced by Glu991 in ScVip1, which adopts different conformations in our apo and substrate-bound crystal structures (Figs. 3c; 8a, b). Replacement of Glu991 with alanine activates ScVip1^PD^ (Fig. 3d, e). In addition, we observed that the ScVip1^PD^ Glu991 to Ala mutant readily accepts 1-InsP_7_ as a substrate in both NMR and malachite green-based phosphatase assays, but remains unable to hydrolyze InsP_6_ (Fig. 8c, d). It has been previously reported that mutation of Asp304 in AppA strongly alters the stereospecificity of the *E. coli* enzyme^60^ Notably, multiple inositol polyphosphate phosphatases that lack strong stereospecificity for InsP_6_ are characterized by an alanine residue at this position (Fig. 8a, b)^61^. We conclude that Glu991, which is conserved among PPIP5Ks from different species (Supplementary Fig. 2) is involved in the selection of the 1,5-InsP_8_ substrate. The inositol ring occupies a different position in our ScVip1^PD^ – 1,5-InsP_8_ complex structure when compared to known phytases (Fig. 8b). We speculate that this is an adaptation to the hydrolysis of a pyrophosphate substrate. The long and uniquely shaped substrate channel may play an additional role in the selection of 1,5-InsP_8_ (Fig. 7c).

**Fig. 8.**
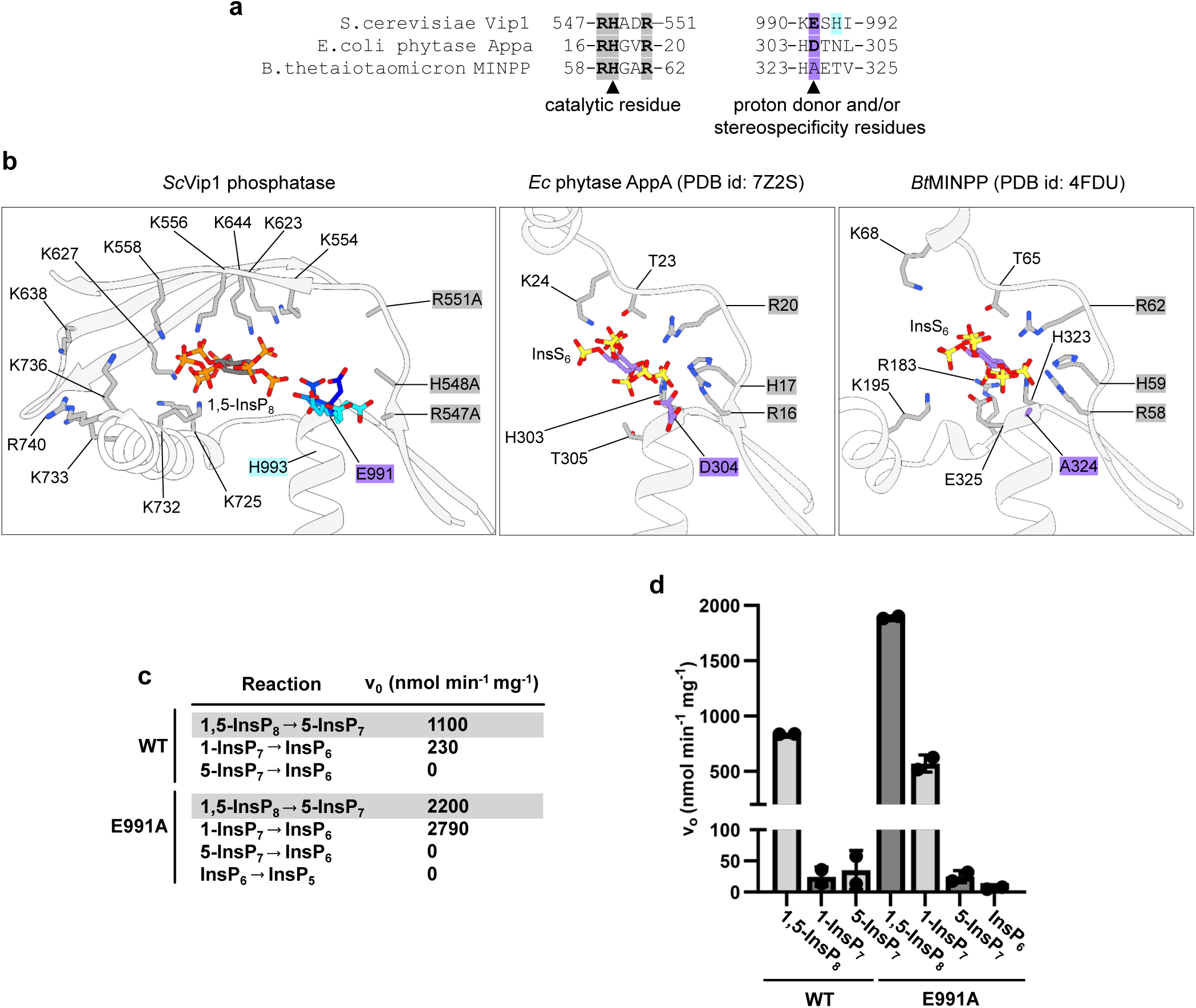
A conserved glutamate controls substrate sterospecificity in ScVip1^PD^. **a** Structure-based sequence alignment of the presumed catalytic motifs in ScVip1^PD^, *E. coli* AppA and the multiple inositol polyphosphate phosphatase MINPP from *Bacteroides thetaiotaomicron*. The RHxxR motif is conserved among all enzymes, while the proton donor aspartate in EcAppA is replaced by Glu991 in ScVip1^PD^ and by an alanine residue in MINPP. **b** Structural comparison of the ScVip1^PD^, EcAppA (PDB-ID 7Z2S^60^) and MINPP (PDB-ID 4FDU^61^) active sites. Glu991 (in bonds representation, coloured from cyan to blue) adopts different conformation in our structures. Asp304 controlling substrate stereospecificity in EcAppA^60^, and Ala324 in the highly promiscuous MINPP enzyme are highlighted in blue. Initial velocities of 1,5-InsP_8_, 1-InsP_7_ and 5-InsP_7_ hydrolysis for wild-type or Glu991Ala versions of ScVip1^PD^, measured by NMR-based (**c**) or malachite green-based (**d**) phosphatase assays.

### Mutation of key motifs inhibit the function of the VIH2 phosphatase domain in planta

We next sought to functionally characterize the substrate binding and catalytic mechanisms of ScVip1^PD^ in yeast. Previous studies in fission yeast revealed a requirement of the PPIP5K Asp1 in ensuring fidelity of chromosome segregation by increasing microtubule stability^35^. Like the fission yeast ortholog, ScVip1 is encoded by a non-essential gene in budding yeast^24^. We analysed the Vip1 genetic interaction network in numerous Synthetic Genetic Array (SGA) screens to identify mutant backgrounds in which Vip1 function becomes critical^62,63^. These studies revealed synthetic sick/lethal interactions between Vip1 and components required to maintain chromosome stability^62^. One such component is Bim1^64^, a conserved +end microtubule binding protein, which in complex with Kar9 forms the cortical microtubule capture site and delays the exit from mitosis when the mitotic spindle is oriented abnormally^65^. We generated a vip1Δbim1Δ mutant strain that is synthetic sick (Supplementary Fig. 9a) and used it for complementation assays. We found that both expression of wild-type full-length ScVip1 and of a mutant targeting the RHxxR motif (ScVip1^R547A-H548A-551A^), which impairs the catalytic function of the phosphatase (Fig. 3d, e), fully restored growth to wild-type like levels (Supplementary Fig. 9a). The ScVip1^E991A^ activating mutant (Fig. 3d,e) also behaved similar to wild type, as did a mutant targeting the outer substrate binding site in the phosphatase domain (ScVip1^K558A-K605A-K732A^) (Supplementary Fig. 9a).

In an alternative approach, we found that overexpression of the isolated kinase domain is lethal for yeast cells grown on low phosphate (Pi) containing media (ScVip1^KD^ in Supplementary Fig. 9b). This effect was not observed when overexpressing either full-length or kinase-dead (ScVip1^KD K414A-D487A^) versions of the enzyme (Supplementary Fig. 9b). In contrast, overexpression of either wild-type or catalytically inactive versions of ScVip1^PD^ was not lethal when yeast were grown on low Pi-containing media (Supplementary Fig. 9b).

Returning to our original plan to characterize the catalytic mechanism of plant PPIP5K phosphatases (see above), we next mapped all residues and sequence motifs characterized in ScVip1 ^PD^ to AtVIH2 from *Arabidopsis thaliana*, which shares 25% sequence identity with ScVip1^PD^ (Fig. 9a; Supplementary Fig. 2). We have previously reported that expression of a phosphatase-dead variant of full-length AtVIH2 (AtVIH2^R372A-H373A^) in Col-0 wild-type background reduced rosette size and intracellular Pi levels^10^. We used these phenotypes to perform structure-function studies with the AtVIH2 phosphatase domain. Constitutive expression of wild-type AtVIH2 had no detectable effect on either rosette area or cellular Pi levels (Fig. 9b-e). Expression of a kinase-dead version of the enzyme (AtVIH2^K219A-D292A^) resulted in hyperaccumulation of cellular Pi, as previously reported (Fig. 9b-3)^10^. Three different mutant combinations targeting the catalytic core of the phosphatase in Arabidopsis VIH2 (compare Fig. 3c-e) were all associated with significantly reduced rosette areas and cellular Pi levels (highlighted in blue in Fig. 9a-e). Mutations targeting the conserved structural Zn^2+^ binding site (compare Fig. 5b-e) were also associated with reduced rosette areas and cellular Pi levels (highlighted in purple in Fig. 9a-e). A mutant in the phosphatase hinge region showed less severe growth phenotypes and only slightly reduced Pi levels (shown in red in Fig. 9a-e), whereas the corresponding mutations in ScVip1^PD^ strongly affected the activity and structural stability of the enzyme *in vitro* (Fig. 5a, c-e). Consistent with the moderate effect of mutations in the α-helical insert contributing to the outer 1,5-InsP_8_ binding site (Fig. 7d-f), the corresponding AtVIH2^R722S-D724A-T726A^ mutant looks similar to the AtVIH2 wild-type expressing control (Fig. 9a-e). Importantly, mutations targeting either the outer or inner GAF domain binding surfaces (compare Fig. 7d-f) both resulted in a severe reduction in rosette area and in low Pi levels (shown in yellow in Fig. 9a-e). In conclusion, our mutational analysis of the AtVIH2 phosphatase domain confirms the importance of the catalytic center of the enzyme and highlights the critical contributions of the outer and inner 1,5-InsP_8_ binding sites in the PPIP5K-specific GAF domain (Fig. 9; Supplementary Fig. 10).

**Fig. 9.**
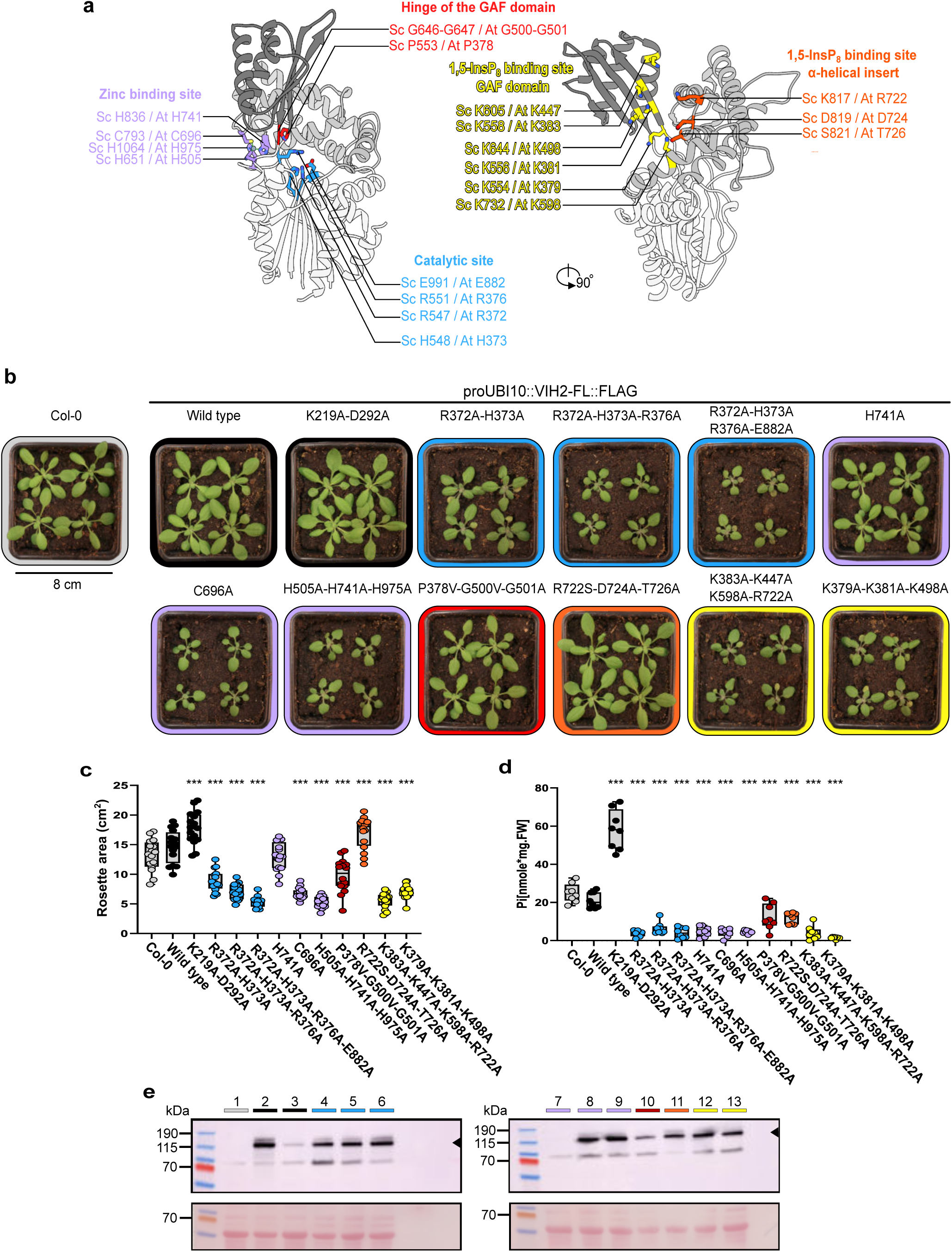
Structure-function analysis of the AtVIH2 phosphatase domain. **a** Ribbon diagram of the ScVip1PD^Δ848-918^ structure, with the histidine phosphatase core in white, the α-helical insert in light gray, and the GAF domain in dark gray. Residues involved in phosphatase catalysis, Zn^2+^ ion coordination, GAF domain movement, and 1,5-InsP_8_ binding are shown in bond representation. Corresponding residues in AtVIH2 are also indicated, illustrating the conserved functional regions. **b** Rosette phenotype of three weeks old Col-0 wild-type plants and lines expressing full-length Flag-tagged versions of AtVIH2 under the control of the UBI10 promoter. **c** Quantification of the rosette area. Box plots show 16 plants per genotype. Multiple comparisons of the genotypes vs. Col-0 were performed according to Dunnett^102^ as implemented in GraphPad prism v10.3.0 (*** p < 0.001, ** p < 0.005, * p < 0.01). **d** Quantification of cellular Pi levels. For each genotype, 8 individual plants were measured using 3 technical replicates. The estimated Pi concentration was normalized by fresh weight. **e** Western blot of Flag-tagged AtVIH2. The theoretical molecular mass of AtVIH2 is ∼118 kDa (indicated by a black arrow; 1:Col-0, 2:AtVIH2^WT^,3:AtVIH2^K219A-D292A^, 4:AtVIH2^R372A-H373A^, 5:AtVIH2^R372A-H373A-R376A^, 6:AtVIH2^R372A-H373A-R376A-E882A^, 7:AtVIH2^H741A^, 8:AtVIH2^C696A^, 9:AtVIH2^H505A-H741A-H975A^, 10:AtVIH2^P378V-G500V-G501A^, 11:AtVIH2^R722S-D724A-T726A^, 12:AtVIH2^K383A-K447A-K598A-R722A^, 13:AtVIH2^K379A-K381A-K498A^.

### The ScVip1 kinase and phosphatase domains are independent enzymatic modules

We expressed and purified full-length SpAsp1 from *E. coli* (Fig. 10a), as previously described^40^, applied the apo enzyme to carbon-coated grids and performed negative-stain electron microscopy (Fig. 10b, see Methods). We calculated a low-resolution envelope from two-dimensional projections of single particles and docked the experimental structures of the sequence-related ScVip1 kinase and phosphatase domains (Fig. 10c, Supplementary Fig. 2). Full-length SpAsp1 adopts a L-shaped conformation (Fig. 10c). The N-terminal kinase and the C-terminal phosphatase domains are perpendicular to each other (Fig. 10c). The exact orientation of the kinase domain is somewhat ambiguous but is reasonably constrained by the size of the 16 amino acid linker (Fig. 10c). In our model, the catalytic centers of the kinase and phosphatase modules are far apart, at least in the conformation adopted by the apo enzyme on the electron microscope grid (Fig. 10c). Consistent with this more open conformation of PPIP5K, the inositol 1-pyrophosphate phosphatase activity of ScVip1^KD-PD^ (residues 186-1107) is comparable to that of ScVip1^PD^ (Fig. 10d). Next, we performed hydrogen/deuterium exchange mass spectrometry experiments on ScVip1^KD-PD^. Addition of 1,5-InsP_8_ to the apo enzyme protects regions from the α-helical insertion domain and the GAF domain (Fig. 10e; Supplementary Fig. 11), further supporting our substrate binding model for ScVip1^PD^ (Fig. 6c), and the proposed dynamics of the GAF domain (Figs. 5a, 7a-c). Addition of an ATP substrate analog reveals additional protected regions in the vicinity of the active site of the kinase domain (Fig. 10e; Supplementary Fig. 11). No changes consistent with the formation of new kinase – phosphatase domain interfaces are observed upon addition of either the phosphatase or kinase substrate (Fig. 10e; Supplementary Fig. 11). Taken together, these experiments suggest that the PPIP5K kinase and phosphatase modules are enzymatically and structurally independent entities.

**Fig. 10.**
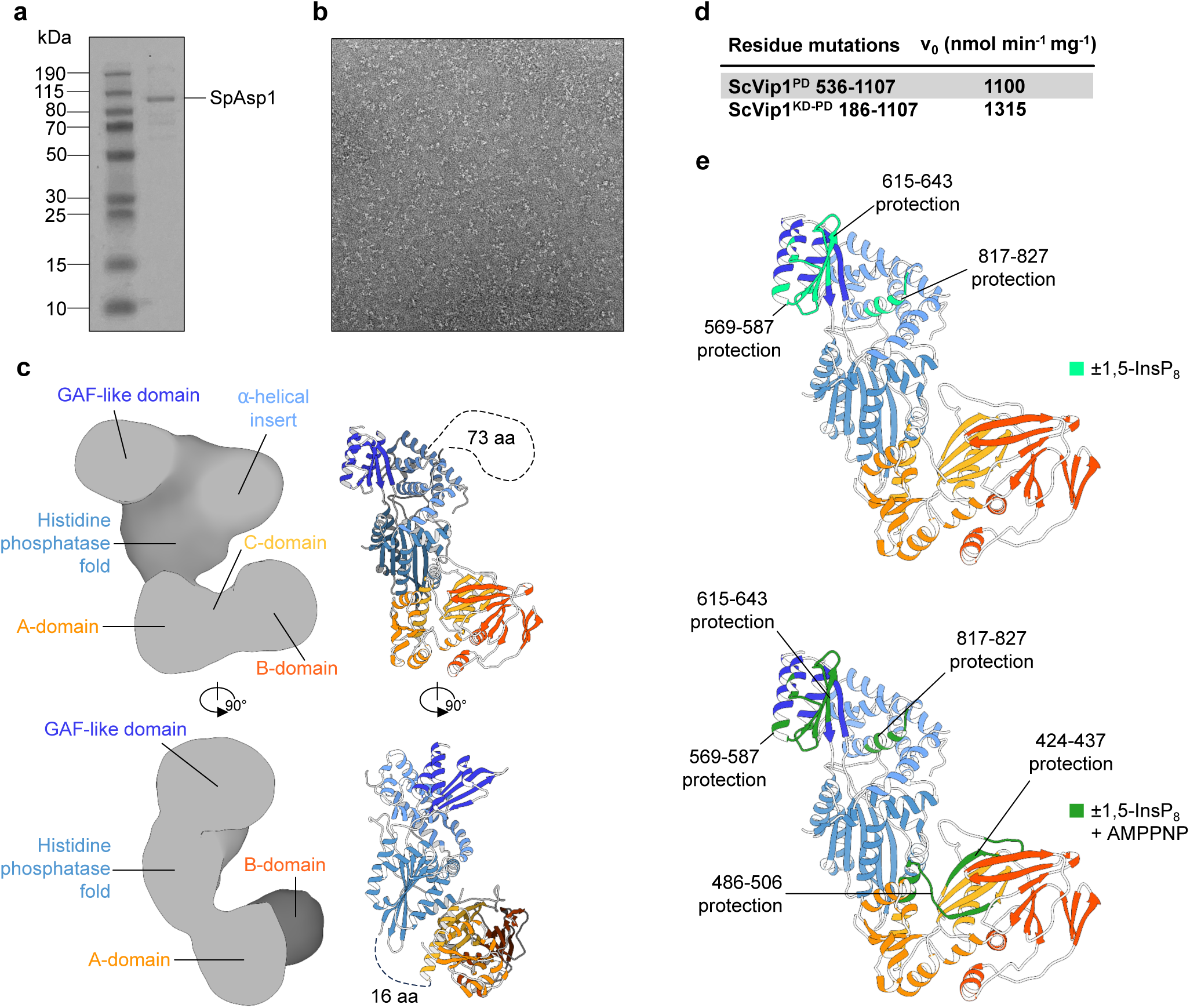
Arrangement of the the kinase and phosphatase domains in a full-length PPIP5K. **a** Coomassie blue-stained SDS-PAGE of purified SpAsp1. **b** A representative micrograph of the negatively stained SpAsp1 used for image processing. **c** Low resolution envelope of SpAsp1 determined by negative stain electron microscopy (left). Individual crystal structures of ScVip1KD (orange ribbon diagram) and ScVip1^PD Δ848-918^ (blue ribbon diagram) were docked into the L-shaped negative stain map of SpAsp1 (right). The linkers truncated in the crystal structures of ScVip1 are potentially visible in the SpAsp1 envelope (black dashed lines). **d** Initial velocities of 1,5-InsP_8_ hydrolysis catalyzed by ScVip1^PD^ or ScVip1^KD-PD^, measured by NMR-based phosphatase assays. **e** Hydrogen-deuterium exchange data mapped on the ScVip1^KD-PD^ model. The protected regions in ScVip1 after incubation with non-hydrolysable 1,5-InsP_8_ (PCP-IP_8_) are shown in light green, the protected regions after incubation with non-hydrolysable 1,5-InsP_8_ (PCP-IP_8_) and ATP (AMPPNP) analogs, respectively are shown in dark green.

## Discussion

PPIP5Ks play a central role in inositol pyrophosphate metabolism and signaling^2^. The enzyme activity, substrate specificity, 3-dimensional structure and catalytic mechanism of the N-terminal kinase domain of PPIP5K are well established^26–28,40,46,66^, but the C-terminal phosphatase domain has only been partially characterized biochemically^25,29,40–42^, and has resisted structural analysis^2^. We were able to overcome this problem by focusing on the phosphatase domain of ScVip1, by using a eukaryotic expression system, by fine-mapping the phosphatase domain boundaries and by engineering of a large unstructured loop (see Methods, Table 1). The structure of ScVip1^PD^ confirms its evolutionary relationship with bacterial and animal histidine acid phosphatases^24^. We found that the active site of ScVip1^PD^ is very similar to that of InsP_6_ hydrolyzing phytases, with both enzymes sharing the structurally conserved RHxxR catalytic motif^24,47^ (Fig. 3c-e).

A second catalytic histidine (His993 in ScVip1, His807 in SpAsp1) was originally defined by sequence comparison with *E. coli* AppA^23,24^, where the corresponding His303 is involved in InsP_6_ substrate binding and in catalysis^56,57^. Structural comparisons of EcAppA and ScVip1^PD^ show that this motif has been incorrectly assigned. His303 in the bacterial phytase corresponds to a poorly conserved lysine residue (Lys990) in ScVip1 (Supplementary Figs. 11, 2). His993 in ScVip1 or His807 in SpAsp1 are not part of the active site but rather map to an α-helical segment buried in the phosphatase domain (Supplementary Fig. 12). Thus, substitution of SpAsp1 H807 with alanine may disrupt the structural integrity of the enzyme rather than affect its catalytic activity, providing an alternative explanation for the reduced enzyme activities and associated phenotypes reported for this mutant variant^35,41,42^. Our *in vitro* (Fig. 3) and *in planta* (Fig. 9) analyses of the substrate binding and catalytic mechanism of the enzyme now provide a reliable set of mutant combinations for engineering phosphatase-dead PPIP5Ks for genetic rescue experiments in various eukaryotes. Our complementation assays in yeast (Supplementary Fig. 9) and in Arabidopsis ^10^ both suggest that the catalytic activity of the PPIP5K phosphatase domain is dispensable for growth. This is likely due to apparent partial redundancy (biochemical and genetical) of PPIP5K phosphatases with other inositol pyrophosphate phosphatases in yeast ^67,68^ and in plants^34^.

A conserved aspartate (Asp304) has been previously reported as the proton donor in *E. coli* phytase^58^ and mutation of Asp304 strongly alters the substrate stereospecificity of this enzyme ^60^. In ScVip1^PD^ this aspartate is replaced by glutamate (Glu991), which is conserved among PPIP5Ks from different species, and which adopts different conformations in our apo and substrate-bound structures (Fig. 8a,b). The ScVip1^E991A^ mutant exhibits an overall increase in enzyme activity and reduced stereospecificity (Fig. 8c,d). We hypothesize that both Asp304 in *E. coli* phytase and Glu991 in ScVip1^PD^ influence the size and charge distribution of the substrate binding site, with mutations increasing the processivity of the enzyme at the expense of substrate selectivity.

Mapping previously identified SpAsp1 missense mutations from a genetic suppressor screen onto our ScVip1^PD^ structure revealed that the His686 (His836 in ScVip1^PD^) to Tyr and Cys643 (Cys793 in ScVip1^PD^) to Tyr mutants target the structural Zn^2+^ binding site in PPIP5K (Fig. 1c). This further highlights the contribution of the Zn^2+^ binding site to the structural integrity of the enzyme^69^ (Supplementary Fig. 7).

In the case of ScVip1^PD^, a *K*_M_ of ∼5 μM has been reported for 1-InsP_7_^25^. Using saturating substrate concentrations, we have derived *K*_M_ values of ∼70 μM and 45 μM for 1-InsP_7_ and 1,5-InsP_8_, respectively, and 1,5-InsP_8_ is the preferred substrate of ScVip1^PD^ *in vitro* (Fig. 3a,b). For comparison, the PP-InsP phosphatase Siw14 from yeast has a *K*_M_ for 5-InsP_7_ of ∼10 μM^70^. The cytosolic concentration of 1,5-InsP_8_ in *S. cerevisiae* has been recently estimated at ∼0.3 μM^5^. However, the local concentration in proximity to the N-terminal Vip1 kinase domain may be higher.

Our structures reveal that the insertion GAF domain is a distinctive structural feature of PPIP5K phosphatases (Fig. 2a). In phytases, the β1-insert maps to the position of the GAF domain. It has been previously demonstrated that this β1-insert undergoes conformational changes upon InsP_6_ binding^56^ (Fig. 2). Our structures reveal that the PPIP5K GAF domain undergoes much larger conformational changes and plays a pivotal role in substrate binding and in channeling of the substrate to the active site (Figs. 5-7). This is further highlighted by the growth phenotypes of mutant versions of AtVIH2 targeting the GAF domain (Fig. 9). Despite the low degree of sequence conservation, substrate binding, catalysis and regulation appear to be well conserved among PPIP5Ks from different organisms (Fig. 9).

In our 1,5-InsP_8_ complex structure, one substrate molecule is bound to the outer GAF domain, the second molecule maps close to the active site (Fig. 6a-c). These structures and our mutational analysis define inositol polyphosphates/pyrophosphates as novel ligand candidates for GAF domains^52^. It is noteworthy that a PP-InsP “capture site” outside the active site has also been identified in the PPIP5K kinase domain ^45^. Orthophosphate is a known inhibitor of human and yeast PPIP5K phosphatase domains, thereby regulating 1,5-InsP_8_ levels and contributing to phosphate homeostasis^10,16,40^. We speculate that the inhibitory effect of orthophosphate in ScVip1^PD^ (Fig. 4a) may be caused by the binding of this metabolite to the highly basic substrate channel, where it may block access of the 1,5-InsP_8_ substrate to the outer substrate binding surface (Fig. 7a,c). We attempted to further characterize this intriguing regulatory mechanism, but ScVip1^PD^ crystals grown in the presence of Pi, sulfate or tungstate did not diffract.

The low-resolution envelope of full-length SpAsp1 and our hydrogen/deuterium exchange mass spectrometry experiments collectively indicate that the PPIP5K kinase and phosphatase domains share only a small interface. The active sites of the kinase and phosphatase domains are situated at a considerable distance from one another, which is incompatible with the hypothesis of direct substrate channeling (Fig. 10c,e). A similar L-shaped arrangement of kinase and phosphatase domains has been previously observed in the crystal structure of fructose-2,6-bisphosphate kinase phosphatase^71^. Consistent with this very open architecture of the enzyme, the phosphatase activity of full-length ScVip1 is not drastically different from that of isolated ScVip1^PD^ (Fig. 10d). However, this appears not to be the case for the kinase activity in human PPIP5K2, where the isolated kinase domain appeared much more active compared to the full-length enzyme^37^. The domain boundaries of the PPIP5K kinase and phosphatase domains suggest the presence of large unstructured N- and C-terminal loops, as well as the occurrence of a large disordered linker region within the α-helical insertion domain the phosphatase module (Fig. 1a,c; Supplementary Fig. 2). These sequence-diverse linker regions may represent protein interaction sites ^37,72^ that could alter the architecture and regulation of the full-length enzyme. Moreover, the linkers are found phosphorylated in yeast (Supplementary Fig. 2), suggesting that additional regulatory mechanisms of PPIP5K remain to be elucidated.

## Methods

### Expression construct cloning

The kinase (residues 186-522, ScVip1^KD^), phosphatase (residues 536-1107, ScVip1^PD^), loop-truncated phosphatase (residues 536-1107, Δ848-918, ScVip1PD^Δ848-918^), and kinase-phosphatase tandem domains (residues 186-1107, ScVip1^KD-PD^) from *Saccharomyces cerevisiae* Vip1 were cloned from synthetic genes (Twist Bioscience) codon-optimized for expression in *Trichoplusia ni* (Tnao) cells (Invitrogen) in vector pBB3, providing tobacco etch virus protease (TEV) cleavable N-terminal 10xHis and twin StrepII affinity tags (Supplementary Table 1). Point mutations in the ScVip1 phosphatase domain were generated by site-directed mutagenesis and subcloned in pBB3 (Supplementary Table 1). For crystallization optimization, the ScVip1^PD Δ848-918^ construct was engineered by replacing the flexible loop with a short Gly-Ser-Ser-Gly linker. The ScVip1^PD Δ848-918^ construct was further optimized for co-crystallization with 1,5-InsP_8_ by replacing Arg547, His548 and Arg551 in the catalytic center with alanines. Full-length Asp1 from *Schizosaccharomyces pombe* was cloned from a synthetic gene codon-optimized for expression in *Escherichia coli* into the pMH-HSSumo vector, providing N-terminal small ubiquitin-like modifier fusion protein (Sumo) containing 7xHis and StrepII affinity tags, as described^40,73^.

### Protein expression and purification

ScVip1^KD^, ScVip1^PD^, ScVip1^PD Δ848-918^, ScVip1PD ^Δ848-918 RHR-AAA^, ScVip1^KD-PD^, and their point-mutated versions were expressed in Tnao cells (Invitrogen) using recombinant baculoviruses^74^. Typically, 10 mL of recombinant P3 baculovirus was used to infect 200 mL of Tnao insect cells growing in SF900 media (Gibco) at a cell density of approximately 2.0 × 10^6^ cells/mL. The cells were incubated for 72 h at 28°C with shaking at 110 rpm. Cells were harvested by centrifugation at 1,000 g for 20 minutes at 4°C when cell viability was 80–85% and GFP fluorescence indicated >50% infection rate. The cell pellets were flash-frozen in liquid N _2_ and stored at -80°C in lysis buffer containing 50 mM Bis-Tris HCl (pH 7.5), 500 mM NaCl, protease inhibitor cocktail tablets (cOmplete EDTA-free; Roche), and DNase I (Roche). For protein purification, cells were thawed and lysed by sonication (Branson Sonifier DS450). The lysate was clarified by ultracentrifugation at 55,000 g for 60 min at 4°C and filtered through 0.45 µm (Durapore Merck) filters. The supernatant was loaded onto a Ni^2+^ affinity column (HisTrap HP 5 mL, Cytiva), washed with 50 mM Bis-Tris HCl (pH 7.5), 500 mM NaCl, 20 mM imidazole, 0.5 mM EDTA, and eluted with wash buffer supplemented with 300 mM imidazole (pH 8.0) directly onto a Strep-Tactin XT 4Flow 5 mL column (IBA). The final elution from the Strep-Tactin column was performed using BXT buffer (IBA) supplemented with NaCl to reach a final concentration of 500 mM. The eluted fractions were incubated with TEV protease for 16 h at 4°C to remove the 10xHis and twin StrepII affinity tags. The cleaved protein was separated from the tags by a second Ni^2+^ affinity chromatography step. Further purification was achieved by size-exclusion chromatrography on a HiLoad Superdex 200 16/600 pg column (Cytiva) equilibrated in 20 mM HEPES (pH 7.5) and 150 mM NaCl. The peak fractions were concentrated to 5-15 mg/mL and immediately used for crystallization experiments or flash-frozen in liquid nitrogen and stored at -80°C for enzymatic assays.

The expression and purification protocol for full-length SpAsp1 was derived from the work of Benjamin et al.^40^. Protein expression was done in *E. coli* BL21 (DE3) RIL cells grown in Terrific Broth containing 50 μg/mL kanamycin, 34 μg/mL chloramphenicol, and 1% (v/v) ethanol. When the culture reached an OD_600nm_ of 0.8, protein expression was induced with 0.5 mM isopropyl β-D-1-thiogalactopyranoside (IPTG) at 18°C for 16 h. Cells were harvested by centrifugation and resuspended in a buffer containing 50 mM Tris-HCl (pH 8.0), 500 mM NaCl, and 10% glycerol, supplemented with a cOmplete protease inhibitor cocktail tablet, DNase I, and 0.5 mg/mL lysozyme (Roth). Cell lysis was achieved by sonication. The lysate was then clarified by centrifugation at 46,500 g for 1 h, and the supernatant was loaded onto a Ni^2+^ affinity column (HisTrap HP 5 mL, Cytiva) equilibrated in 50 mM Tris-HCl (pH 8.0), 500 mM NaCl, and 10% (v/v) glycerol. After a washing step with the same buffer, SpAsp1 was eluted with buffer supplemented with 250 mM imidazole (pH 8.0). The 7xHis-StrepII-SUMO tag was cleaved by treatment with Ulp1 protease during an overnight incubation at 4°C^75^. The tag-free SpAsp1 protein was separated from the fusion tag by a second Ni^2+^ affinity chromatography step. SpAsp1 was then diluted into 50 mM Tris-HCl (pH 8.0) and 10% (v/v) glycerol to reach a final NaCl concentration of 50 mM and loaded onto a HiTrap Heparin HP 5 mL column (Cytiva) equilibrated with 50 mM Tris-HCl (pH 8.0), 50 mM NaCl, and 10% (v/v) glycerol. SpAsp1 was eluted against a gradient of 50 mM Tris-HCl (pH 8.0), 1 M NaCl, and 10% (v/v) glycerol. Finally, SpAsp1 was applied to a HiLoad Superdex 200 16/600 pg column (Cytiva) equilibrated in 20 mM HEPES (pH 7.5) and 150 mM NaCl. The peak fractions containing SpAsp1 were concentrated and immediately used for negative stain electron microscopy.

### Crystallization and data collection

ScVip1^KD^ (8.0 mg/mL in 20 mM Hepes pH 7.5, 150 mM NaCl) was incubated with 5 mM ADP and 20 mM MgCl_2_ for 1 h on ice prior to crystallization. Crystals developed in hanging drops composed of 1.0 μL of protein solution and 1.0 μL of crystallization buffer (23 % [v/v] PEG 3,350, 0.1 M citric acid / BIS-Tris propane pH 6.4) suspended over 1.0 mL of the latter as reservoir solution. A complete dataset to 1.18 Å resolution was collected at beam line X06DA of the Swiss Light Source, Villigen, Switzerland (Table 1). Triclinic crystals of apo ScVip1^PD^ developed in hanging drops composed of 1.5 μL of protein solution (15 mg/mL in 20 mM Hepes pH 7.4, 150 mM NaCl) and 1.5 μL of crystallization buffer (20 % [v/v] PEG 3,350, 0.2 M NH_4_NO_3_). Crystals were derivatized for 18 h in crystallization buffer supplemented with 0.1 mM K_2_PtCl_4_ and subsequently snap-frozen in liquid N_2_ after serial transfer in crystallization buffer supplemented with 20 % (v/v) glycerol as cryoprotectant. A single anomalous diffraction (SAD) dataset at the ‘white line’ L3 edge was collected at beam line X06DA (Table 1). Native crystals of ScVip1 ^PD^ developed in 20% v/v (+/-)-2-Methyl-2,4-pentanediol, 0.1 M Tris pH 8.0. Crystals were cryoprotected by addition of glycerol to a final concentration of 20% (v/v) and a complete dataset with apparent orthorhombic symmetry was collected at beam line X06DA to 3.05 Å resolution (Table 1). Crystals of the engineered ScVip1^PDΔ848-918^ construct (10 mg/mL in 20 mM Hepes pH7.5, 150 mM NaCl, 0.5 mM (CH_3_CO_2_)_2_Zn) developed in 2 μL drops containing 10 % (w/v) polyvinylpyrrolidone K15, 5 mM CoCl_2_, 0.1M Tris pH 7.0 as crystallization buffer and 0.1 μL additive solution (5mM 13:0 Lyso PC aka 1-tridecanoyl-2-hydroxy-sn-glycero-3-phosphocholine) using microseeding protocols. Crystals were cryoprotected by serial transfer into crystallization buffer containing glyerol to a final concentration of 30% (v/v) and snap frozen in liquid N_2_. Data collection at beam line X06DA yielded a complete dataset at 3.4 Å resolution (Table 1). The ScVip1^PD Δ848-918 RHR-AAA^ – 1,5-InsP_8_ complex crystallized in sitting drops containing 0.25 μL protein solution (12 mg/mL in 20 mM Hepes 7.5, 150 mM NaCl, 2.5 mM 1,5-InsP_8_) and 0.25 μL of crystallization buffer (Morpheus II screen condition H2, 32.5% v/v Precipitant Mix 6: 25% w/v PEG 4000, 40% w/v 1,2,6-Hexanetriol; 0.04M Polyamines: 0.01M Spermine tetrahydrochloride, 0.01M Spermidine trihydrochloride, 0.01M 1,4-Diaminobutane dihydrochloride, 0.01M DL-Ornithine monohydrochloride; 0.1M Buffer System 4: pH6.5 MOPSO, Bis-Tris)^76^. Crystals were cryoprotected by serial transfer in crystallization buffer supplemented with ethylene glycol to a final concentration of 25 % (v/v) and diffracted up to 2.36 Å resolution (Table 1). Data processing was done with XDS^77^ (version June 30, 2023).

### Crystallographic structure solution and refinement

The structure of ScVip1^KD^ was solved by molecular replacement method using the program PHASER ^78^ and the isolated kinase domain of HsPPIP5K2 as search model (https://www.rcsb.org/ PDB-ID 3T9A). The solution comprises a monomer in the asymmetric unit. The structure was completed in alternating cycles of manual model building in COOT^79^ and restrained refinement with anisotropic atomic displacement parameters as implemented in PHENIX.REFINE^80^. The structure of ScVip1^PD^ was solved by molecular replacement combined with single anomalous diffraction (MR-SAD) using a low redundancy SAD dataset collected from a K_2_PtCl_2_-derivatized crystal and a fragment of the *Francisella tularensis* histidine acid phosphatase core domain (PDB-ID 3it3) in the program PHASER-EP^81^. The resulting solution comprised four molecules in the asymmetric unit, four Pt^2+^ and four Zn^2+^ ions (Table 1). The starting phases and model were used for non-crystallographic symmetry (NCS) density modification in PHENIX.RESOLVE^82^ and the resulting 3.5 Å electron density map was sufficient to build a nearly complete poly-alanine model that was used to determine the structure of ScVip1^PD^ apo by molecular replacement. ScVip1^PD^ apo crystals had an apparent *C* 2 2 2 symmetry, however no packing solution could be identified with PHASER^78^ and analysis with PHENIX.XTRIAGE suggested an abnormal distribution of intensities. The structure was subsequently solved in space group *P* 1 and input into the program ZANUDA^83^ for space group validation. ZANUDA returned a solution in space group *P* 2_1_, with four molecules in the asymmetric unit (Table 1). The a and c axes are almost identical, with a twin fraction of ∼0.5, similar to previous reports^84,85^. The structure was completed in alternating cycles of manual model building and restrained NCS refinement with twin operator l, -k, h in PHENIX.REFINE^80^. A large, apparently disordered loop (residues 851-918) was identified in this structure and subsequently replaced by a short Gly-Ser-Ser-Gly linker. The structures of ScVip1^PDΔ848-918^ and ScVip1^PD Δ848-918 RHR-AAA^ were solved by molecular replacement using the ScVip1 apo structure as search model and refined with PHENIX.REFINE^80^ (Table 1). The stereochemistry of the refined models was assessed with PHENIX.MOLPROBITY^86^. Structural diagrams were generated with CHIMERA^87^.

### NMR-based phosphatase assay

ScVip1 phosphatase assays were performed as previously described^10^. Reactions contained 100 µM of the respective [^13^C_6_]-labeled PP-InsP in 20 mM HEPES pH 7.0, 150 mM NaCl, 0.2 mg/mL BSA and D_2_O to a total volume of 600 µL. Reactions were supplemented with sodium phosphate, sodium sulfate or sodium nitrate, as indicated. Reaction mixtures were pre-incubated at 37°C and the reaction was started by adding the respective amount of enzyme (ScVip1^PD^ (WT): 10 nM for 1-InsP_7_ and 1,5-InsP_8_, 700 nM for 5-InsP_7_; 1.5 µM R547A-H548A-R551A; 1.25 µM R547A; 1.5 µM H548A; 1.5 µM R551A; 10 nM Δ848-918; 1.5 µM K554A; E991A: 2 µM for InsP_6_, 20 nM for 1-InsP_7_, 10 nM for 1,5-InsP_8_; 1 µM P553V-G646V-G647A; 10 nM C793A; 10 nM H651A; 350 nM K558A-K605A-K732A-K817A; 1 µM K554A-K556A-K644A; 10 nM K817S-D819A-S821A; 50 nM E576A-K623A-Q625S-K627A). Turnover of PP-InsPs was monitored continuously at 37°C using a nuclear magnetic resonance spectroscopy (NMR) pseudo-2D spin-echo difference experiment^88^. Individual NMR spectra were recorded every 86 s. The relative intensity changes of the C2 peaks of the respective PP-InsPs as a function of reaction time were used for quantification^10,88^. Data analysis was carried out with GraphPad PRISM 5. Trend lines were added to the raw data, following either a linear regression model or the first derivative of the equation of a one phase decay model, normalized by the enzyme’s mass concentration.

In order to determine the substrate preference of ScVip1^PD^, reaction mixtures containing 20 mM HEPES pH 7.0, 150 mM NaCl, 0.2 mg/mL BSA and 100 µM of the respective [^13^C_6_]-labeled (PP-)InsP were prepared and pre-incubated at 37°C. ScVip1^PD^ was added subsequently (100 nM for 1-InsP_7_ and 1,5-InsP_8_, 1 µM for InsP_6_ and 5-InsP_7_) and the reactions were incubated at 37°C for 3 h. As a control, H_2_O was added instead of the enzyme. Reactions were quenched by heating to 95°C for 5 min. The samples were centrifuged at 15,000 g for 5 min and the supernatant was analyzed by a ^1^H-^13^C-BIRD-HMQC 2D NMR as described^88^.

### Malachite green-based phosphatase assay

The phosphatase activity of ScVip1^PD^ was determined using the Malachite Green Phosphate Assay Kit (Sigma-Aldrich). This assay detects the release of inorganic phosphate catalyzed by ScVip1PD from substrates such as 1,5-InsP_8_, 1-InsP_7_, 5-InsP_7_ or InsP_6_ during the phosphatase reaction^89,90^. For the assay, 100 nM of ScVip1^PD^ was incubated at 37°C for 2,4, and 8 min in 100 µL reaction mixtures containing 20 mM Hepes (pH 7.5), 150 mM NaCl and varying concentrations of substrates (1,5-InsP_8_ ranging from 8 µM to 700 µM; 1-InsP_7_ ranging from 1 µM to 2000 µM, unless indicated otherwise). The reaction was quenched by mixing 40 µL of the reaction mixture with 10 µL of malachite green reagent. The color formation was measured after 25 min on a plate reader (Tecan Sparc) at 620 nm. When precipitation of the reagent occurred at high phosphate concentrations (>100 µM), the reaction mixture was 2-8 times diluted in buffer prior quenching. The quantity of released inorganic phosphatase was estimated using the standard curve of OD ^620 nm^ versus phosphate standard concentrations. Blanks OD_620 nm_ were measured without enzyme at each time point to discard effect of non-enzymatic hydrolysis of PP-InsPs or the eventual presence of inorganic phosphate in the buffer.

### Thermal shift assay

Thermal shift assays were conducted with 25 to 100 µM of wild-type or mutant ScVip1^PD^ in 20 mM Hepes (pH 7.5), 150 mM NaCl, and 8fold dilution of SYPRO Orange dye (Thermo Fisher Scientific). Protein samples were heated with an increasing gradient of 0.05°C/s from 25 to 99°C and melting curves were recorded using a QuantStudio 5 Real-Time PCR System system (Thermo Fisher Scientific).

### Analytical size-exclusion chromatography

Gel filtration experiments were performed using a Superdex 200 Increase 10/300 GL column (GE Healthcare) pre-equilibrated in 20 mM Hepes pH 7.5, 150 mM NaCl. 200 μl of the respective protein was loaded sequentially onto the column, and elution at 0.75 ml/min was monitored by ultraviolet absorbance at 280 nm. Peak fractions were analyzed by SDS–PAGE gel electrophoresis.

### Yeast strains and plasmids

The *Saccharomyces cerevisiae* strains used in this study are listed in Supplementary Table 2. Genomic disruptions at genomic loci were performed as previously reported^91^. Preparation of media, yeast transformations, and genetic manipulations were performed according to established protocols. The plasmids used in this study are listed in Supplementary Table 2, primers used for site-directed mutagenesis in Supplementary Table 1. All recombinant DNA techniques were performed according to established procedures using Escherichia coli TOP10 cells for cloning and plasmid propagation. Gene mutations were generated with QuickChange site-directed mutagenesis kit (Agilent Technologies, Santa Clara, CA, USA). All cloned DNA fragments and mutagenized plasmids were verified by Sanger-sequencing (Microsynth, AG).

### Molecular cloning and generation of stable transgenic A. thaliana lines

The Golden Gate system was used to generate plasmids for Arabidopsis transformation^92^. Mutant versions of the AtVIH2 coding sequence were synthesized (Twist bio-science, San Francisco,CA) (Supplementary Table 1). The different mutants were then cloned into the level one (L1) vector (Supplementary Table 1). To construct the binary vector L1 vectors containing the AtUBI10 promoter (proUBI10), the corresponding AtVIH2 coding sequence, a FLAG tag, the Nos terminator and Fast Red as a fluorescent marker for the seed coat (Supplementary Table 2). The plasmids were transformed into *A. tumefaciens* strain GV3101. Flower dipping was then used to transform 5-week-old plants (Col-0) ^93^. T1 plants were selected by red fluorescence using a Nikon SMZ18 stereomicroscope equipped with an RFP-B filter. Lines with single T-DNA insertions were selected by the segregation analysis. All experiments were performed using stable T3 generation lines (Supplementary Table 3).

### Plant material, seed sterilization, and plant growth conditions

Seeds were sterilized using the chloride gas protocol. Seeds were then sown on ^½^MS media containing 1% (w/v) sucrose, MES (0.5 g/L), pH 5.7 and agar (8g/L). 120 mm square Petri dishes were incubated for 48h at 4°C in the dark and transferred to a growth chamber at 22 °C with a 16h/8h light/dark cycle under a fluorescent light source. 7d old seedlings were transferred to soil in 8 cm pots and grown as indicated.

### Image acquisition and rosette area quantification

To evaluate the rosette phenotypes of the different T3 lines, 16 plants (in 4 pots) were analyzed. After 2 weeks in soil, top images were recorded and the rosette area was manually segmented in Fiji^94^. Specifically, images were converted to 8 bit followed by manual selection of individual plants. Contrast was adjusted, and the rosette was selected using the threshold. The scale (240 px to 1.5 cm) was set before quantification. Finally, the area of the segmented image was determined using the Particle Quantify function as implemented in Fiji.

### Determination of cellular Pi concentration

For Pi quantification 3-weeks-old plants were analyzed (8 independent plants per sample). One cotyledonous leaf per plant was harvested, the fresh weight was recorded and the leaf was transferred to an Eppendorf tube (1.5 ml) containing 500µl miliQ H_2_O. The tubes were then frozen in liquid nitrogen and boiled at 80 °C for 5 min. This step was repeated two times. Measurement was performed using a colorimetric molybdate assay^89^. The master solution contains contained 72 µL of ammonium molybdate solution (0.0044 % [w/v] of ammonium molybdate tetrahydrate, 0.23 % [v/v] of 18 M H _2_SO_4_), 16 µL of 10 % (w/v) acetic acid and 12 µL of miliQ H_2_O. For each reaction, 20 µl of sample were added in triplicate to a 96-well plate and incubated with 100 µl of working solution for 1 h at 37 °C. Absorbance was measured at 820 nm in a Spark plate reader (Tecan). The concentration was determined according to the standard curve with a concentration range of 1, 0.5, 0.2, 0.1 and 0 mM Na_2_HPO_4_.

### Western blotting

Proteins were extracted from 3-weeks-old plants. About 100 mg of sample were collected in 2 ml Eppendorf tubes and frozen in liquid nitrogen. The tissue was homogenized in a tissue lyzer (MM400, Retsch). Then, 300 µl of 1x PBS buffer supplemented with the plant-specific protease inhibitor cocktail (PE0230, Merck) was added to each tube and incubated for 30 min at 4 °C in an orbital shaker (Intelli-Mixer™ RM-2M, ELMI). The samples were then centrifuged at 10,000 g for 20 min at 4 °C. Then, 150 µl of the supernatant was transferred in a new tube containing 30 µl of SDS loading buffer and incubated at 95 °C for 5 min. A 10% BIS-Tris acrylamide SDS-PAGE was run with the MES running buffer. Proteins were transferred to a 0.2 µ PVDF membrane (IB34002, Thermo Scientific™) using a semi-dry protocol (iBlot3, Thermo Scientific). Membranes were blotted with TBS-Tween (0.1%)-milk (5%) for 1 h at room temperature. For FLAG detection, membranes were incubated with a monoclonal Anti-FLAG-M2-Peroxidase (HRP) antibody (A8592, Merck) using a dilution of 1:5000. Detection was performed using SuperSignal West Femto Maximum Sensitivity Substrate (34095, Thermo Scientific).

### Electron microscopy and image processing

The full-length SpAsp1 envelope was analyzed using negative-stain electron microscopy. A total of 5 μl of monomeric peak fractions from a gel filtration run (concentration 0.01 mg per mL) were applied to glow-discharged, carbon-coated copper grids (400 mesh, Electron Microscopy Sciences). After incubation for 1 min, excess liquid was blotted off. The grids were then sequentially passed through one drop of buffer (20 mM Hepes, pH 7.5, and 150 mM NaCl) and two drops of 2% (w/v) uranyl acetate solution, with the sample incubated in the last drop for 1 min before blotting and air drying. A total of 1,565 micrographs were recorded at the DCI-Geneva (cryoGEnic) electron microscopy platform using a Talos L120C microscope operating at 120 kV and equipped with a Falcon II direct electron detector. Data acquisition was automated using EPU software (Thermo Fisher Scientific) with a pixel size of 1.531 Å and a total electron dose of 52 e-/Å². Data processing was performed using CryoSPARC (version 4.1)^95^. Initially, contrast transfer function (CTF) parameters were estimated using patch-based CTF estimation. Micrographs with a CTF fit better than 10 Å resolution were selected, resulting in 1,497 micrographs. Manual picking of 177 SpAsp1 particles, which were extracted in boxes of 250 × 250 pixels, were used to train the blob picker tuner. After this process, 239 exposures were rejected due to containing either too few or excessively high numbers of picked particles, leaving a particle stack of 270,221 particles (averaging 220 particles per micrograph) with defocus values between 0.3 and 1.3 µm. Two rounds of reference-free 2D class averaging were employed to further refine the dataset and remove the stain artifacts, resulting in 107,485 particles. An ab initio reconstruction with six classes was followed by 2D classification and non-uniform refinement for each class. The best 3D reconstruction, representing full-length SpAsp1, comprised 11,837 particles.

### Hydrogen/deuterium exchange mass spectrometry

HDX-MS experiments were performed at the UniGe Protein Platform (University of Geneva, Switzerland) following an established protocol with minimal modifications^96^. Details of reaction conditions and all data are presented in Supplementary Fig. 11. HDX reactions were performed in 50 μl volumes with a final protein concentration of 2.1 μM of ScVip1 and a 100-fold molar excess of AMP-PNP and / or PCP-IP_8_ ^97^. Briefly, 107 picomoles of protein were pre-incubated with ligands for 1 h on ice in a final volume of 7 μl. The deuterium on-exchange reaction was initiated by adding 43 µl of D_2_O exchange buffer (10 mM Tris pH 8 / 150 mM NaCl in D_2_O) to the protein-ligand mixture. Reactions were carried-out at room temperature for 2 incubation times (30 s, 300 s) and terminated by the sequential addition of 20 μl of ice-cold quench buffer (4 M Gdn-HCl / 1 M NaCl / 0.1 M NaH_2_PO_4_ pH 2.5 / 1 % (v/v) formic acid). Samples were immediately frozen in liquid nitrogen and stored at -80 °C for up to two weeks. All experiments were repeated in triplicate. To quantify deuterium uptake into the protein, samples were thawed and injected in a UPLC system immersed in ice with 0.1 % FA as liquid phase. The protein was digested via two immobilized pepsin columns (Thermo Scientific #23131), and peptides were collected onto a VanGuard precolumn trap (Waters). The trap was then eluted, and peptides were separated with a C18, 300Å, 1.7 μm particle size Fortis Bio 100 x 2.1 mm column over a gradient of 8 – 30 % buffer C over 20 min at 150 μl/ min (buffer B: 0.1 % [v/v] formic acid; buffer C: 100 % acetonitrile). Mass spectra were acquired on an Orbitrap Velos Pro (Thermo), for ions from 400 to 2200 m/z using an electrospray ionization source operated at 300 °C, 5 kV of ion spray voltage. Peptides were identified by data-dependent acquisition of a non-deuterated sample after MS/ MS and data were analyzed by Mascot. All peptides analysed are shown in Supplementary Fig. 11. Deuterium incorporation levels were quantified using HD examiner software version 3.3 (Sierra Analytics), and quality of every peptide was checked manually. Results are presented as percentage of maximal deuteration compared to theoretical maximal deuteration level. Changes in deuteration level between two states were considered significant if >7% and >0.5 Da and p< 0.02 (unpaired t-test) for a single deuteration time. Raw values of deuterium in corporation for each peptides in all conditions are listed in Supplementary Table 4.

## Data availability

Crystallographic coordinates and associated structure factors have been deposited with the Protein Data Bank (http://rcsb.org) doi:10.2210/pdb9gr8/pdb (ScVip1^KD^ – ADP, PDB-ID 8GR8), doi:10.2210/pdb9grh/pdb (ScVip1^PD^ – apo, PDB-ID 9GRH), doi:10.2210/pdb9grn/pdb (ScVip1^PDΔ848-918^ – apo, PDB-ID 9GRN) and doi:10.2210/pdb9gro/pdb (ScVip1^PD Δ848-918 RHR-AAA^ – 1,5-InsP_8_, PDB-ID 9GRO). The mass spectrometry proteomics data have been deposited to the ProteomeXchange Consortium via the PRIDE partner repository^98^ with dataset identifier PXD056020 (doi: 10.6019/PXD056020).

## Acknowlegements

We thank P. Rieu and A. Cargenato for critical reading of the manuscript. This work was supported by Sinergia grant CRSII5_209412 from the Swiss National Science Foundation (to D.F., V.G.P. and M.H.) and by Howard Hughes Medical Institute International Research Scholar Award 55008733 (to M.H.).

## Author contributions

P.R. and K.L. expressed and purified proteins, K.L. and P.R. crystallized proteins and solved and refined structures with the help of M.H.. P.R. and S.M.B. performed enzyme assays. P.R. performed thermal shift assays and size-exclusion chromatography. D.P-C. and V.G.P. designed and performed yeast experiments. F.R. performed plant genetic experiments, Pi measurements and western blotting. P.R. performed and analyzed negative stain electron microscopy experiments. K.L. and O.V. performed and analyzed HDX-MS experiments. P.R. and M.H. wrote the manuscript. All authors edited the manuscript and approved the final document.

## Figure legends

**Supplementary Fig. 1.**
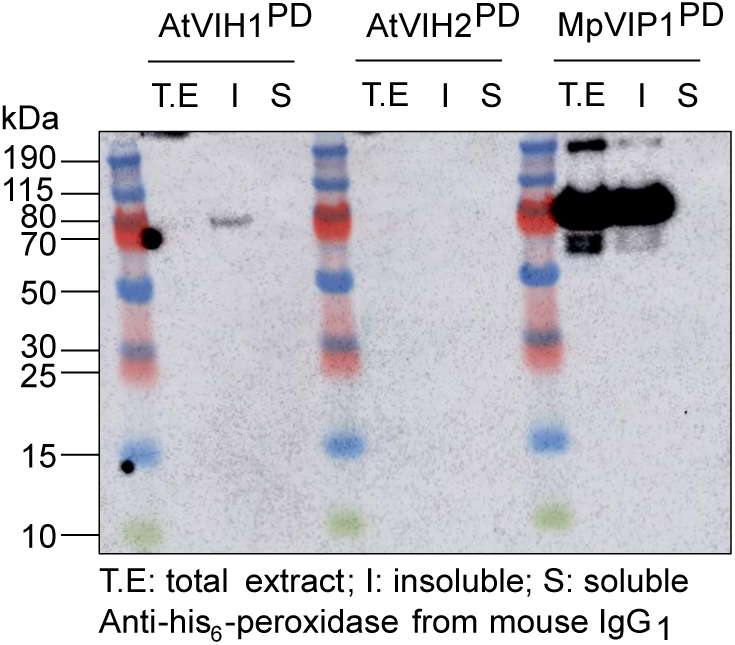
Expression of AtVIH1^PD^, AtVIH2^PD^ and MpVIP1^PD^ phosphatase domains in baculovirus-infected Tnao insect cells. Western blot analysis showing the expression level and solubility of different PPIP5K phosphatase domain (PD) constructs from different plant species. The constructs analyzed are *Arabidopsis thaliana* (*At)* VIH1^PD^ (residues 351-1010, Uniprot ID Q84WW3), VIH2^PD^ (residues 351-1046, Uniprot ID F4J8C6), and *Marchantia polymorpha* (*Mp)* VIP1^PD^ (residues 382-1041 Uniprot ID Mp8g06840).

**Supplementary Fig. 2.**
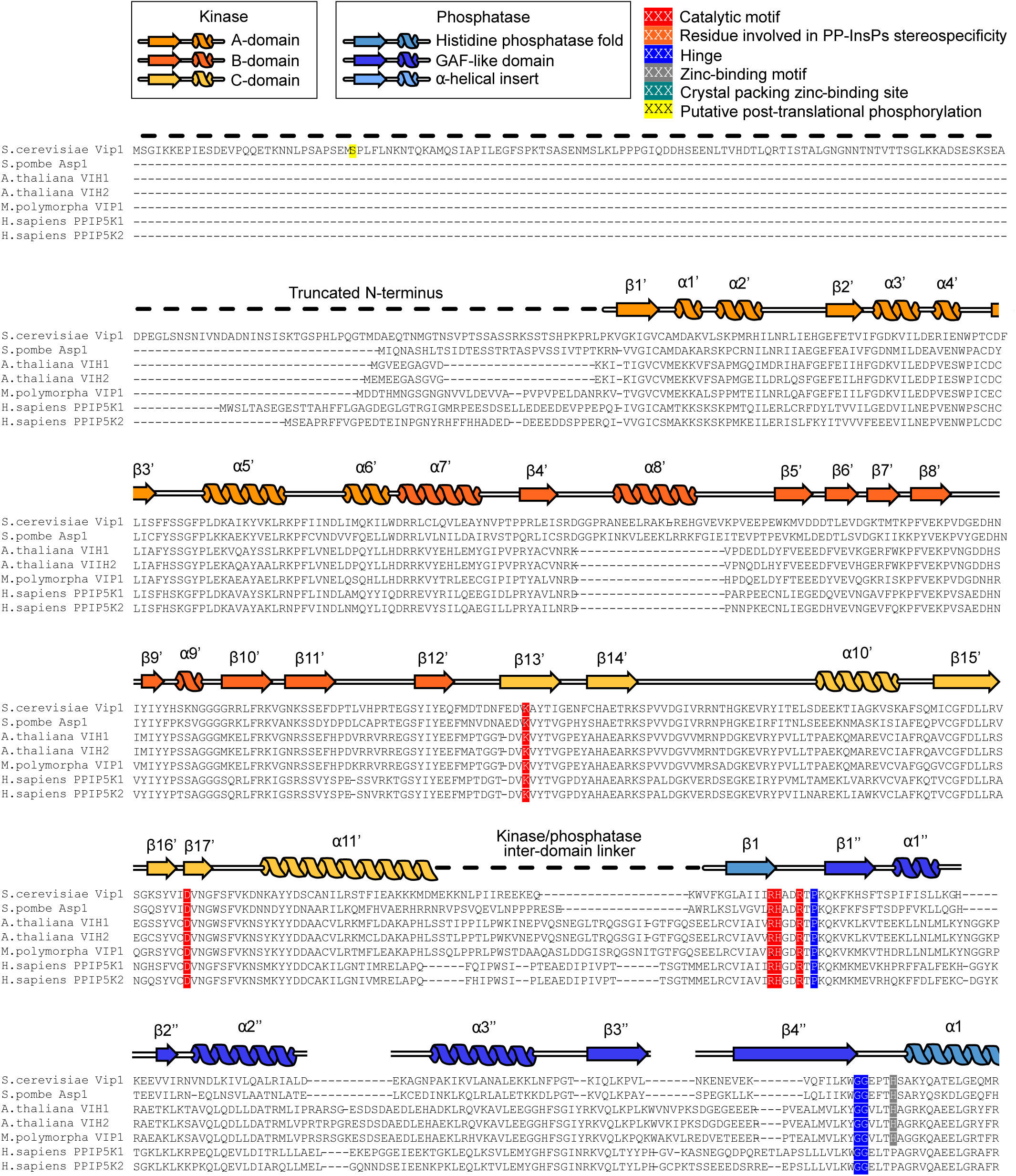
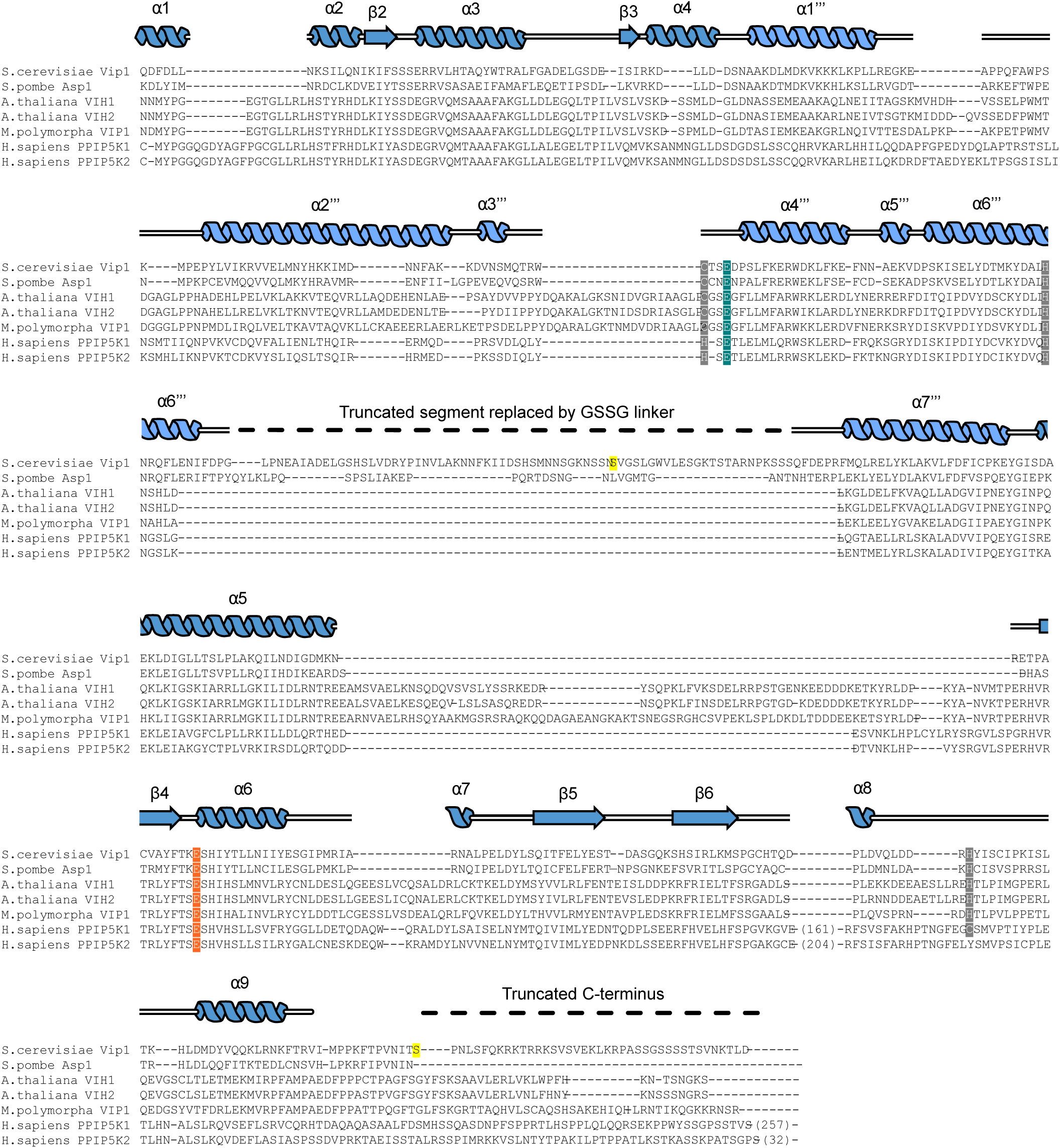
Multiple sequence alignment of PPIP5K phosphatase domains. Multiple-sequence alignment of S. *cerevisiae* Vip1 (Uniprot ID Q06685) with *Schizosaccharomyces pombe* Asp1 (Uniprot ID O74429), *Arabidopsis thaliana* VIH1 and VIH2 (Uniprot ID Q84WW3 and F4J8C6, respectively), *Marchantia polyorpha* VIP1 (Uniprot ID Mp8g06840), and *Homo sapiens* PPIP5K1 and PPIP5K2 (Q6PFW1 and O43314, respectively). Secondary structure elements are colored according to different domains as indicated on top. Functionally important conserved residues are highlighted. Dashed lines indicate regions of the protein truncated in the engineered constructs of ScVip1 kinase domain (residues 186-522) and ScVip1 phosphatase domain (residues 536-1107, Δ848-918) used for crystallization.

**Supplementary Fig. 3.**
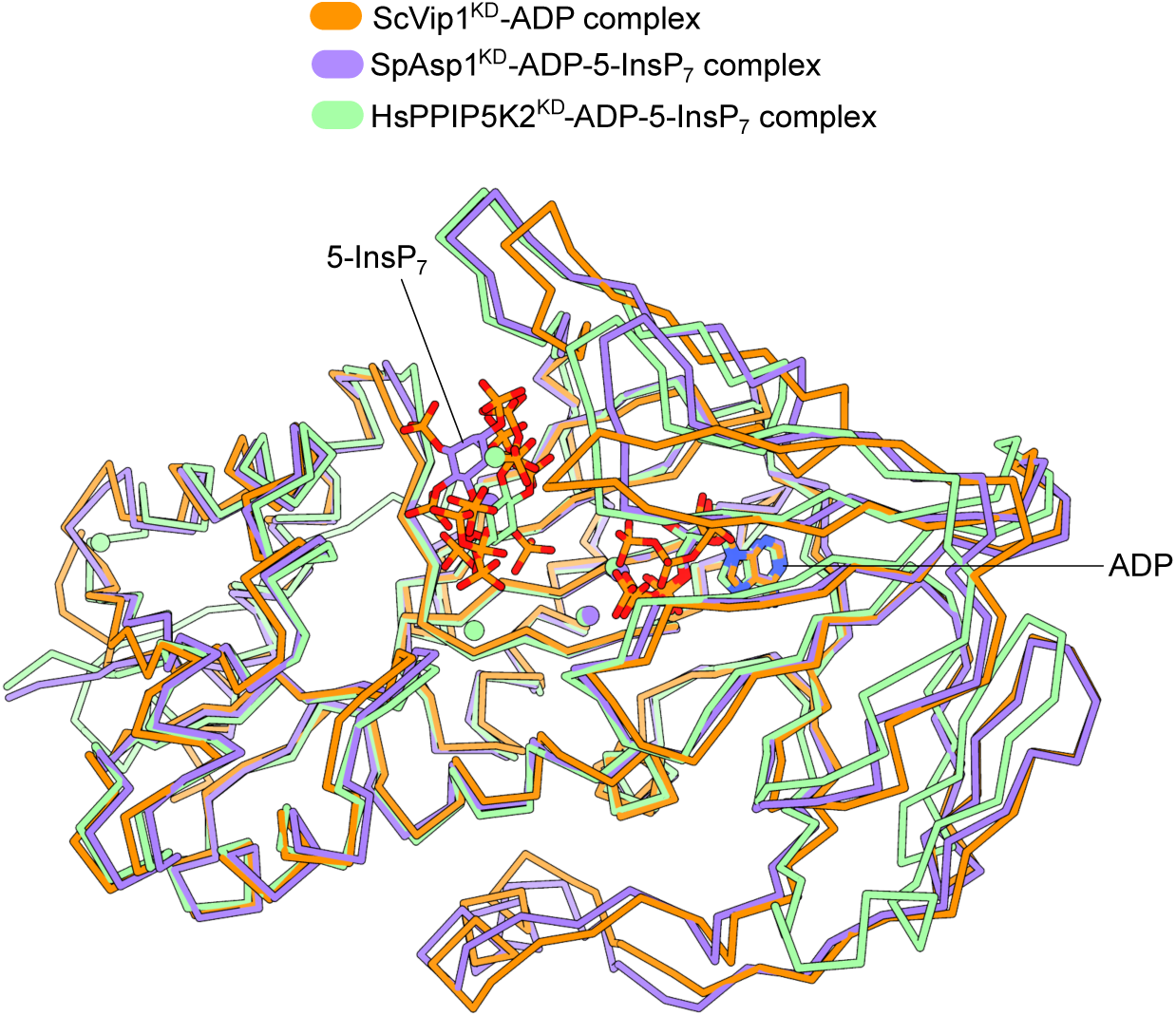
ScVip1 harbors a structurally conserved PPIP5K kinase domain. Structural comparison of different PPIP5K kinase domain structures. Shown are C_α_ traces of the *Saccharomyces cerevisiae* Vip1 kinase domain (in orange, bound to ADP in bonds representation), *Homo sapiens* PPIP5K2 (PDB-ID 3T9E^27^, in green and bound to ADP and 5-InsP_7_) and *Schizosaccharomyces pombe* Asp1 (PDB-ID 8E1H^46^, in purple). The kinase structures align with a r.m.s.d of ∼1.5 Å comparing 310 C_α_ atoms.

**Supplementary Figure 4.**
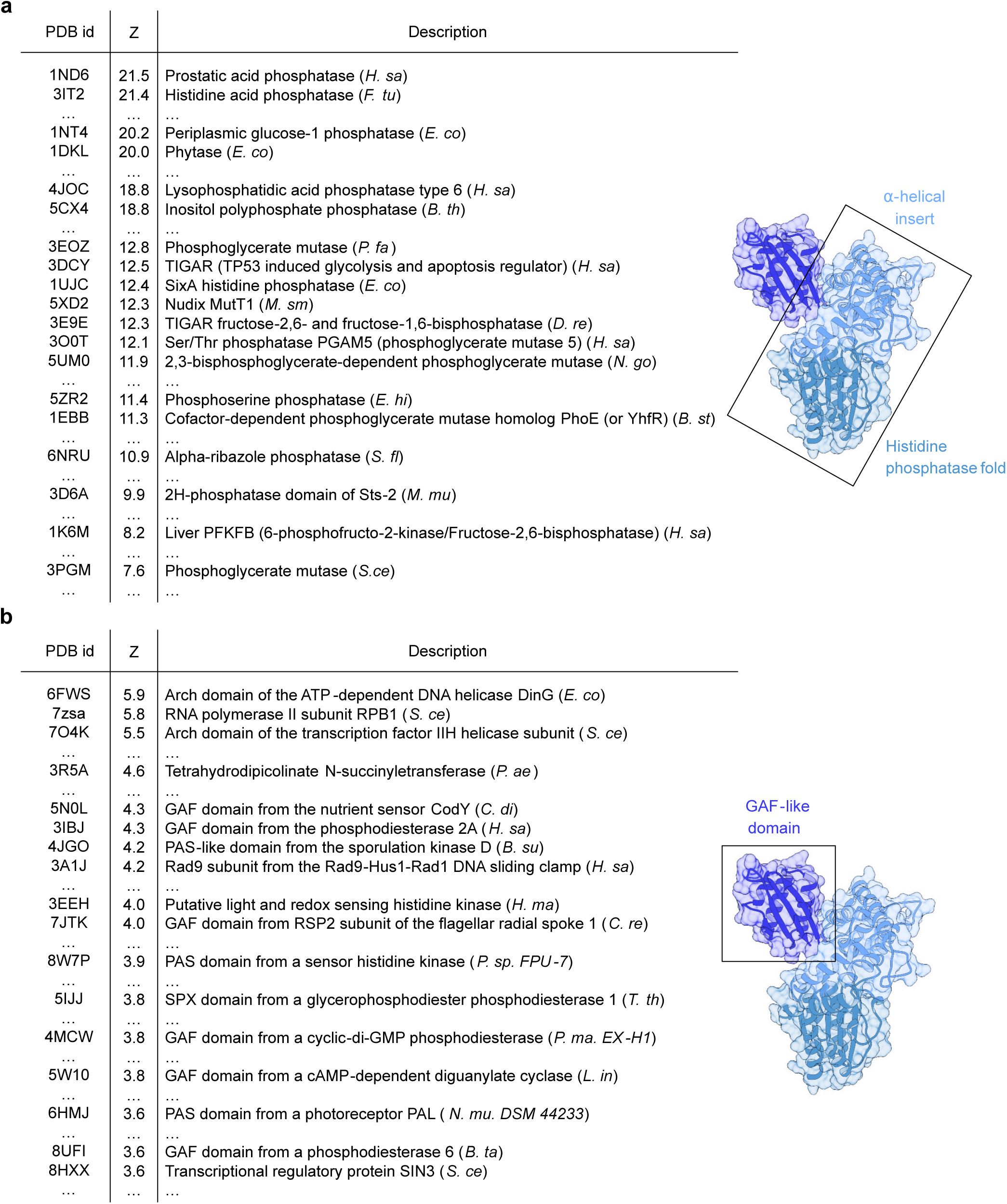
ScVip1^PD^ is structurally related to histidine acid phosphatases and contains a GAF domain. Top twenty hits from the DALI web server (http://ekhidna2.biocenter.helsinki.fi/dali/) using **a** the histidine acid phosphatase core containing the α-helical insertion domain (residues 536-553; 647-1094), or **b** the isolated GAF domain in ScVip1^PD^ (residues 554-646) as search model against the Protein Data Bank.

**Supplementary Fig. 5.**
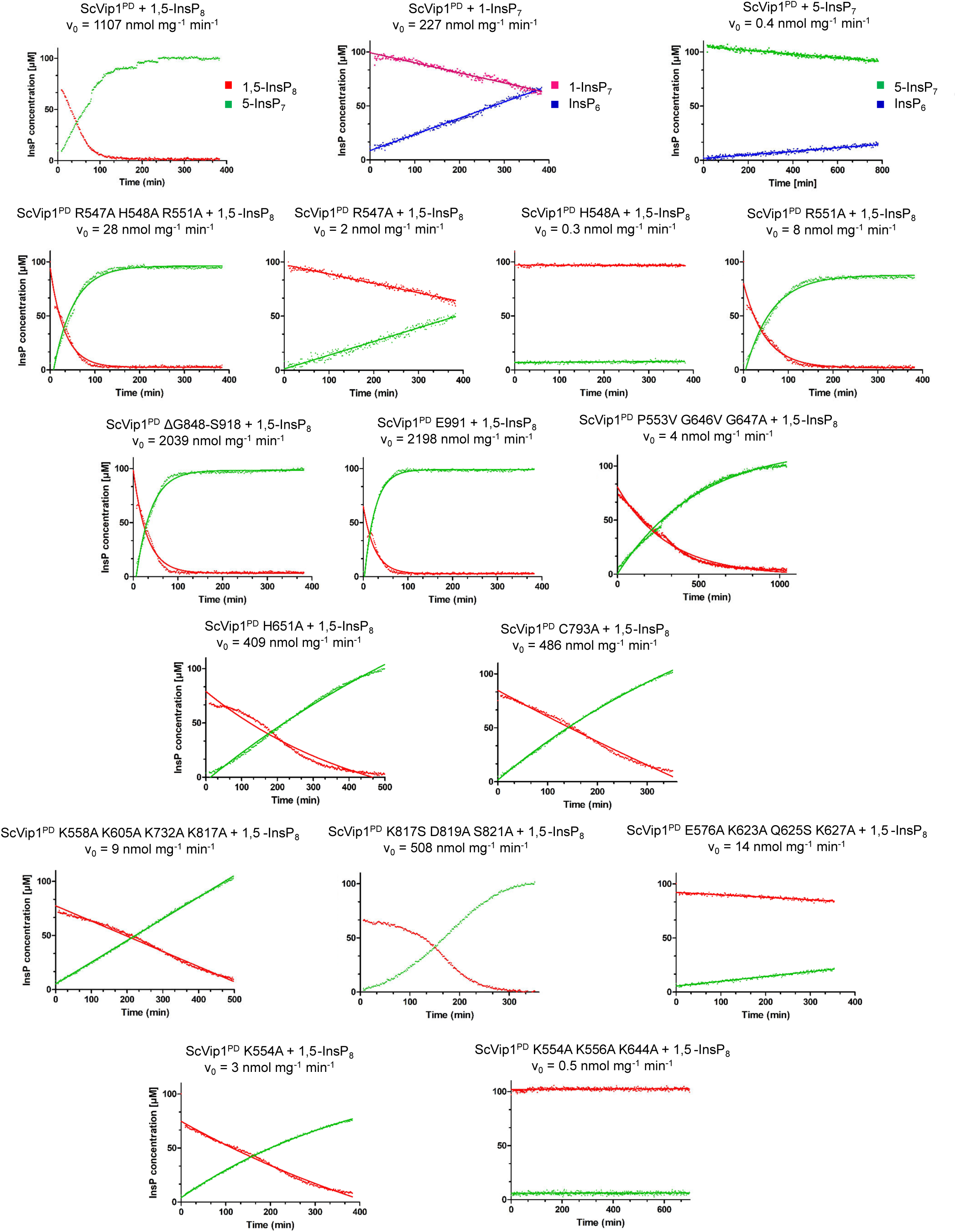
NMR-based enzyme assays – raw data. NMR time course experiments of PP-InsP hydrolysis catalyzed by ScVip1^PD^ and respective mutant constructs using 100µM of [^13^C_6_]-labeled PP-InsP substrates. Concentrations of enzyme were varied based on the respective construct and substrate. PP-InsP turnover was measured using a pseudo-2D spin-echo difference nuclear magnetic resonance spectroscopy experiment. Quantification was based on the relative intensities of the C2 signals of the respective (PP-)InsP species.

**Supplementary Fig. 6.**
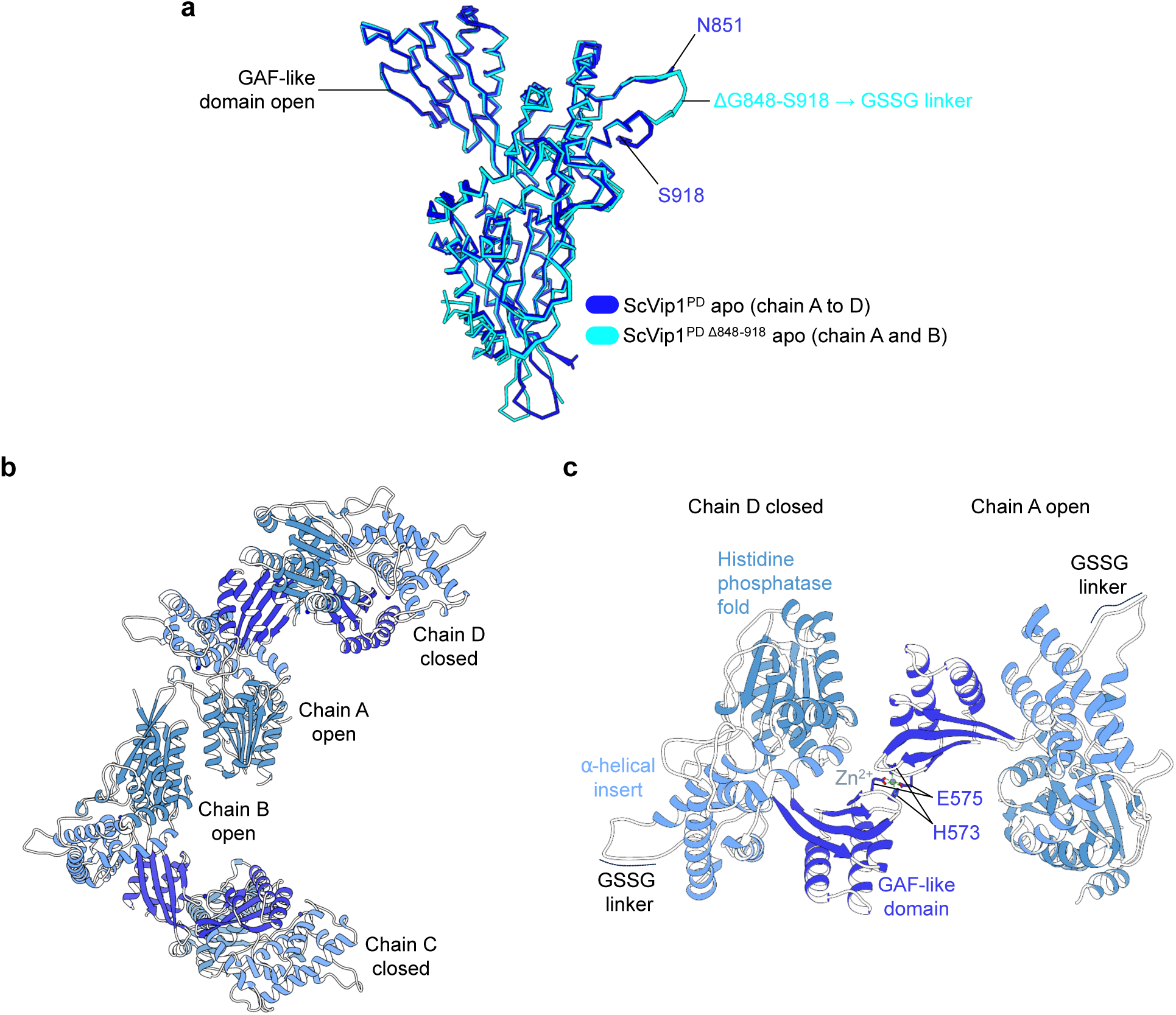
Arrangement of GAF domains in the ScVip1^PD^ and ScVip1^PDΔ848-918^ apo structures. **a** Structural superposition of chains A (shown as C_α_ traces) from the ScVip1^PD^ (in blue) and ScVip1^PDΔ848-918^ (in cyan) apo crystal forms reveal the GAF domain in an open conformation (r.m.s.d is ∼0.5 Å comparing 457 corresponding C_α_ atoms). The asymmetric unit of ScVip1^PD^ apo contains four molecules, all of which present the GAF domain in the open conformation. **b** ScVip1^PDΔ848-918^ crystals also contain four molecules in the asymmetric unit with chains A and B in the open and chains C and D in the closed conformation, respectively. Shown are ribbon diagrams, colors as in Fig. 1b. **c** A second Zn^2+^ binding site in ScVip1^PDΔ848-918^ is formed along a pseudo 2-fold axis involved two neighboring GAF domains. This second binding site likely represent a crystallization artifact. The crystallization condition contained an excess of (CH_3_CO_2_)_2_Zn and the residues involved in Zn^2+^ binding are not conserved among other PPIP5Ks (His573, Glu575, compare Supplementary Fig. 2).

**Supplementary Fig. 7.**
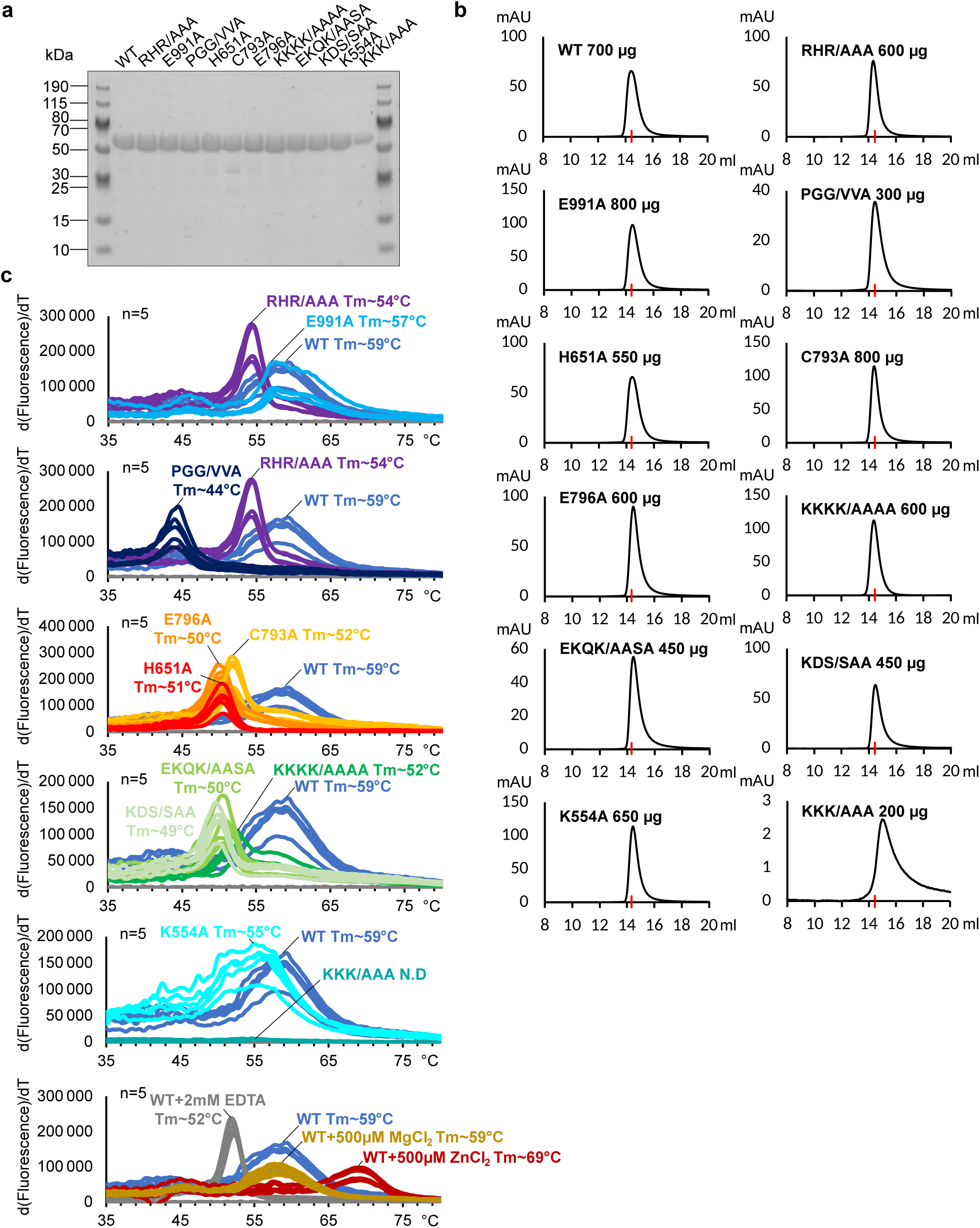
Structural integrity profiling of *Sc*Vip1^PD^ mutant proteins. **a** Coomassie blue-stained SDS-PAGE analysis of wild-type and mutant versions of ScVip1^PD^ (theoretical molecular weight is ∼65.5 kDa). **b** Analytical size-exclusion chromatography of wild-type and mutant versions of ScVip1PD. Shown are A_280nm_ absorption traces as a function of column elution volume (in mL). The red arrow indicates the characteristic elution volume of ScVip1^PD^, which behaves as a monomer in solution. **c** Melting temperatures of wild-type (blue traces, n=5) and mutant versions of ScVip1^PD^ derived from thermal shift assays (N.D. not determined). Note that addition of divalent metal chelators such as EDTA reduces the thermal stability of the enzyme. Addition of ZnCl_2_ but not MgCl_2_ increases ScVip1^PD^ stability, likely by targeting the structural Zn^2+^ binding site.

**Supplementary Fig. 8.**
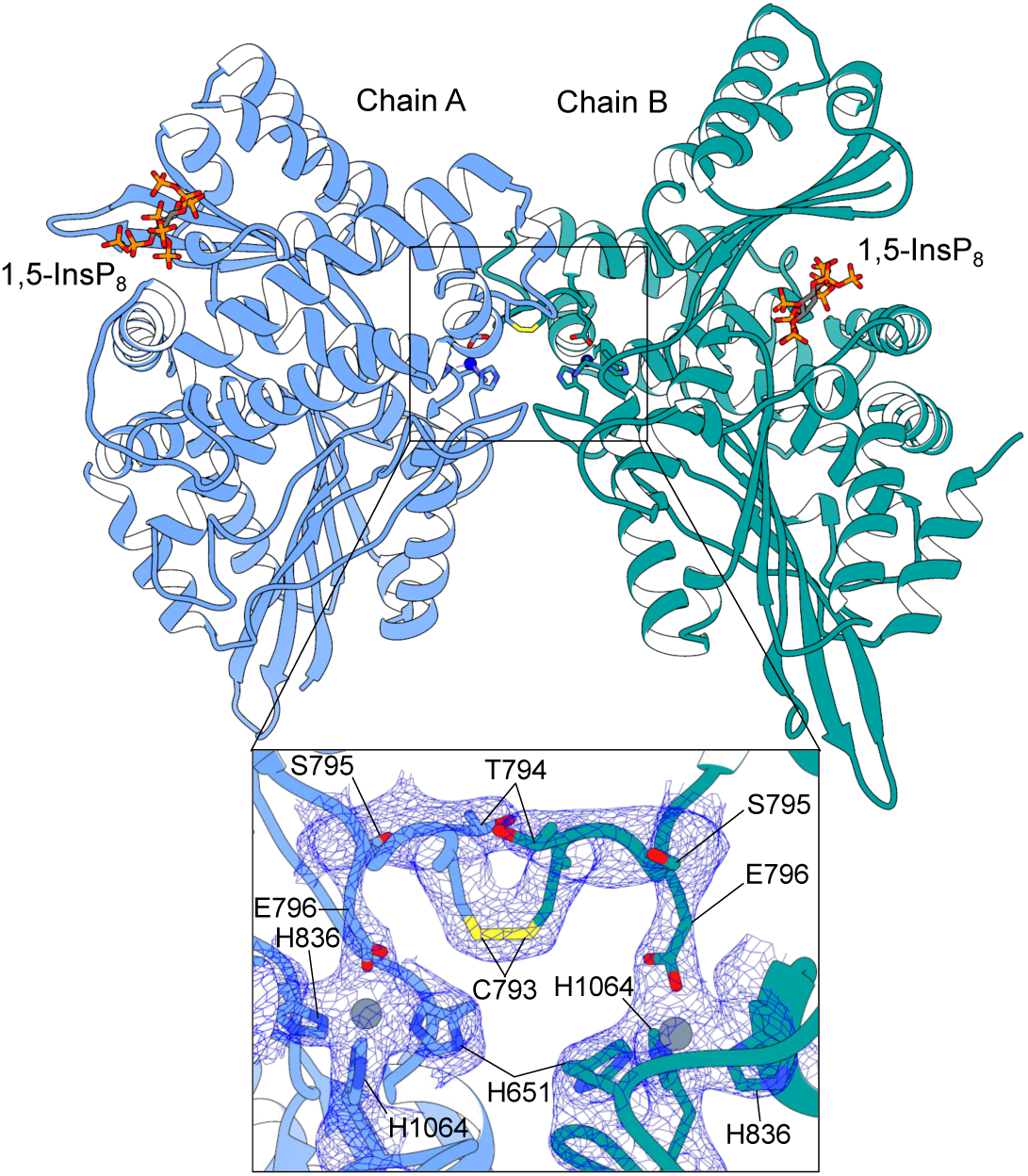
An intermolecular disulfide bridge stabilizes the crystallographic dimer in the *Sc*Vip1^PD Δ848-918 RHR-AAA^ – 1,5-InsP_8_ complex structure. Ribbon diagram of a crystallographic domain-swapped dimer in the asymmetric unit of our ScVip1^PD Δ848-918 RHR-AAA^ crystals (chain A is shown in blue, chain B in green, 1,5-InsP_8_ molecules are depicted in bonds representation). The inlet provides a detailed view of the Zn^2+^ binding site, with the Zn^2+^ ions shown as a sphere (in gray) and the coordinating residues depicted in bonds representation. The view includes a 2 (*F_o_ – F_c_*) electron density map contoured at 1 σ (blue mesh). Cys793 is involved in an intermolecular disulfide bridge between ScVip1^PD Δ848-918 RHR-AAA^ chain A (in blue) and chain B (in green). In the crystal structure of the apo ScVip1^PD Δ848-918^ (compare Fig. 1c), Cys793 replaces Glu796 and coordinates the Zn^2+^ ion.

**Supplementary Fig. 9.**
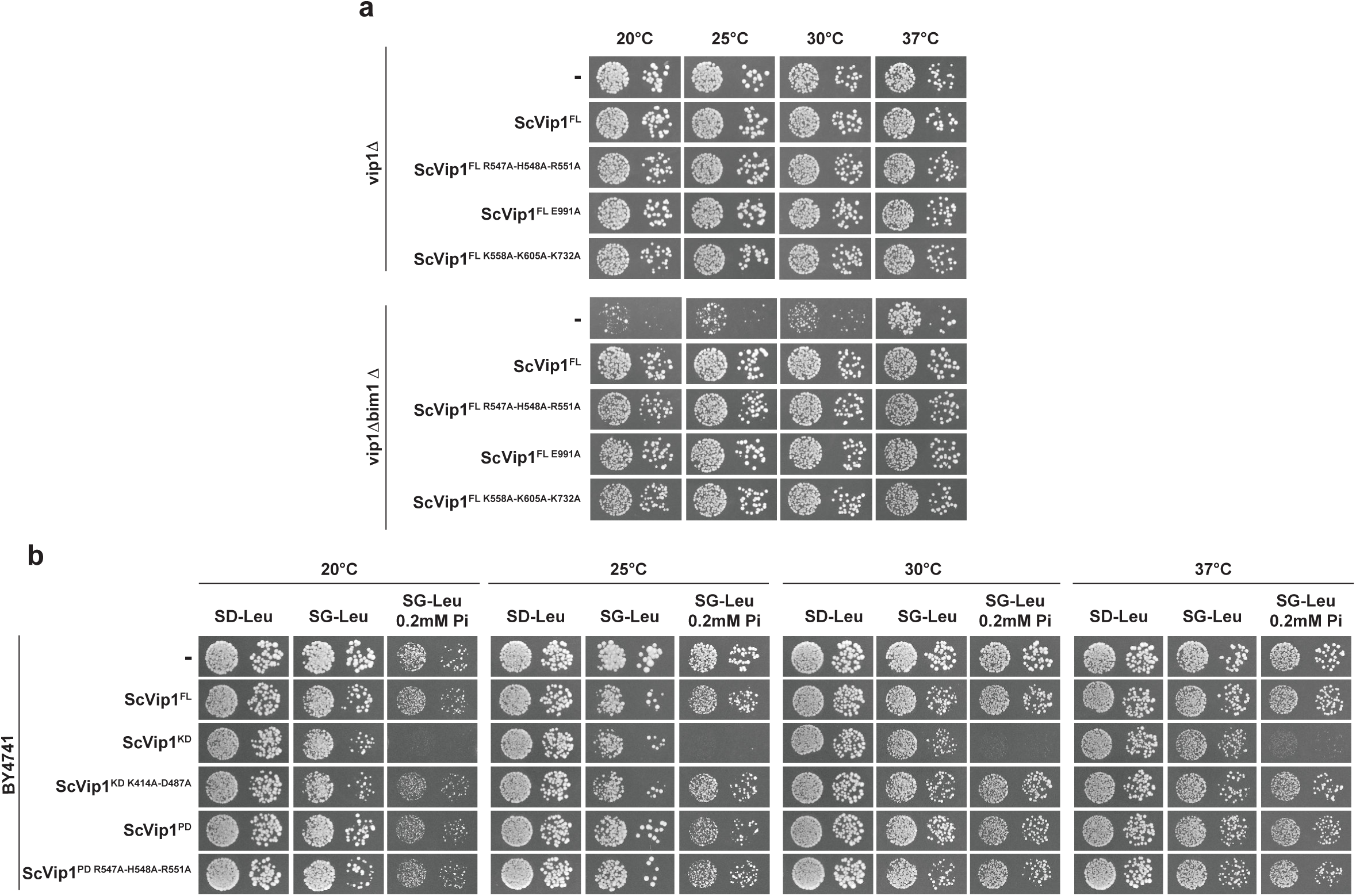
Genetic studies of Vip1 kinase and phosphatase domains in budding yeast. **a** The depicted yeast vip1Δ and vip1Δbim1Δ strains expressing empty vector (-), ScVip1^FL^ (full-length) or ScVip1^FL^ mutants were spotted in 10-fold serial dilutions on glucose- and galactose-containing medium at standard and low concentrations of potassium phosphate (0.2 mM Pi) and incubated at the indicated temperatures for 2–11 days. **b** BY4741 strain expressing empty vector (-), ScVip1^FL^, ScVip1^KD^ (186-524), ScVip1^PD^ (536-1107) or point mutants (as indicated) were spotted in 10-fold serial dilutions on glucose-, galactose-containing medium at standard and low concentrations of potassium phosphate (0.2 mM Pi) and incubated at the indicated temperatures for 2–11 days.

**Supplementary Fig. 10.**
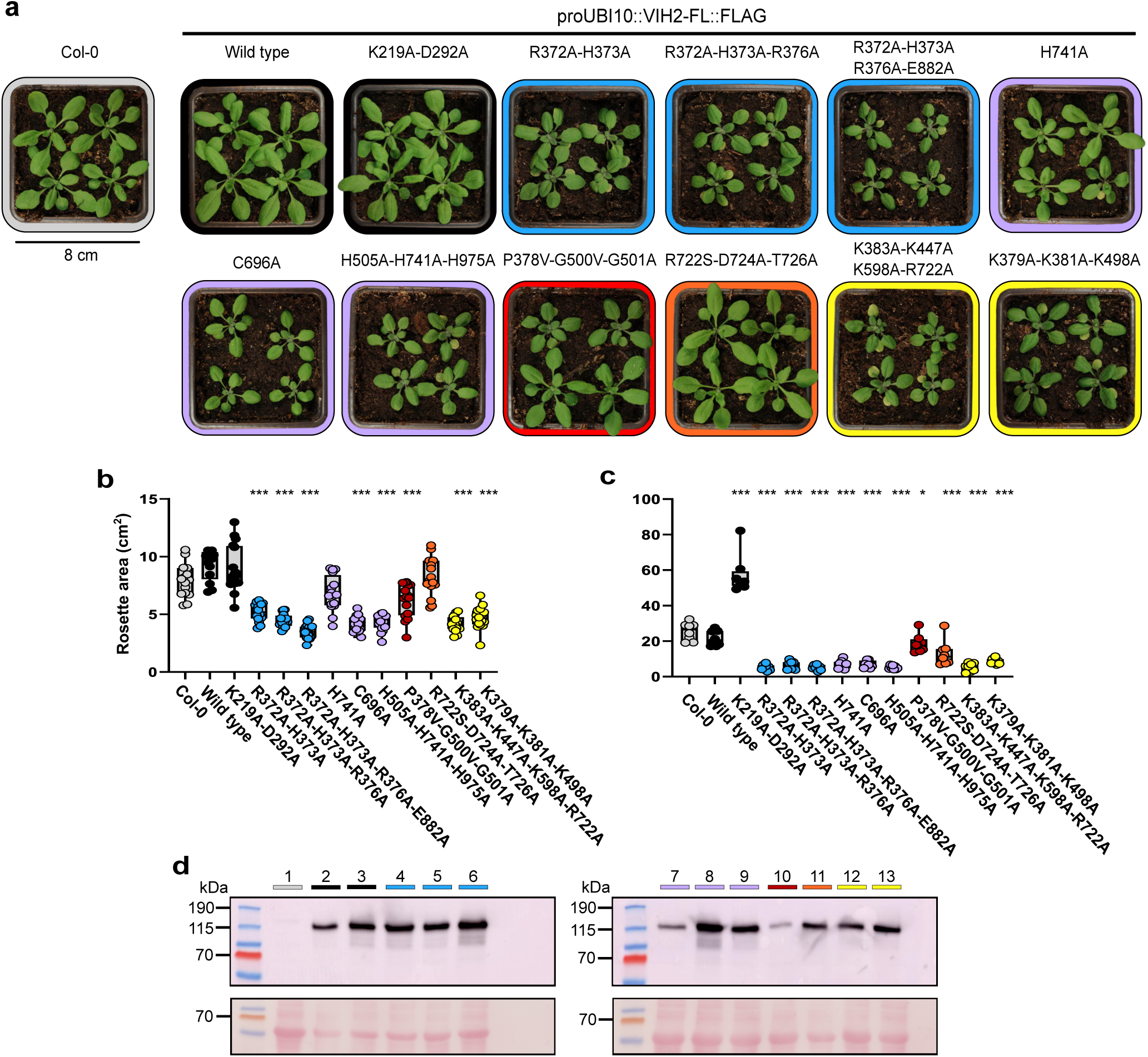
Structure-function analysis of the AtVIH2 phosphatase domain – independent replication experiment. **a** Rosette phenotype of three weeks old Col-0 wild-type plants and lines expressing full-length Flag-tagged versions of AtVIH2 under the control of the UBI10 promoter. **b** Quantification of the rosette area. Box plots show 16 plants per genotype. Multiple comparisons of the genotypes vs. Col-0 were performed according to Dunnett^102^ as implemented in GraphPad prism v10.3.0 (*** p < 0.001, ** p < 0.005, * p < 0.01). **c** Quantification of cellular Pi levels. For each genotype, 8 individual plants were measured using 3 technical replicates. The estimated Pi concentration was normalized by fresh weight. **d** Western blot of Flag-tagged AtVIH2. The theoretical molecular mass of AtVIH2 is ∼118 kDa (indicated by a black arrow; 1:Col-0, 2:AtVIH2^WT^,3:AtVIH2^K219A-D292A^, 4:AtVIH2^R372A-H373A^, 5:AtVIH2^R372A-H373A-R376A^, 6:AtVIH2^R372A-H373A-R376A-E882A^, 7:AtVIH2^H741A^, 8:AtVIH2^C696A^, 9:AtVIH2^H505A-H741A-H975A^, 10:AtVIH2^P378V-G500V-G501A^, 11:AtVIH2^R722S-D724A-T726A^, 12:AtVIH2^K383A-K447A-K598A-R722A^, 13:AtVIH2^K379A-K381A-K498A^.

**Supplementary Fig. 11.**
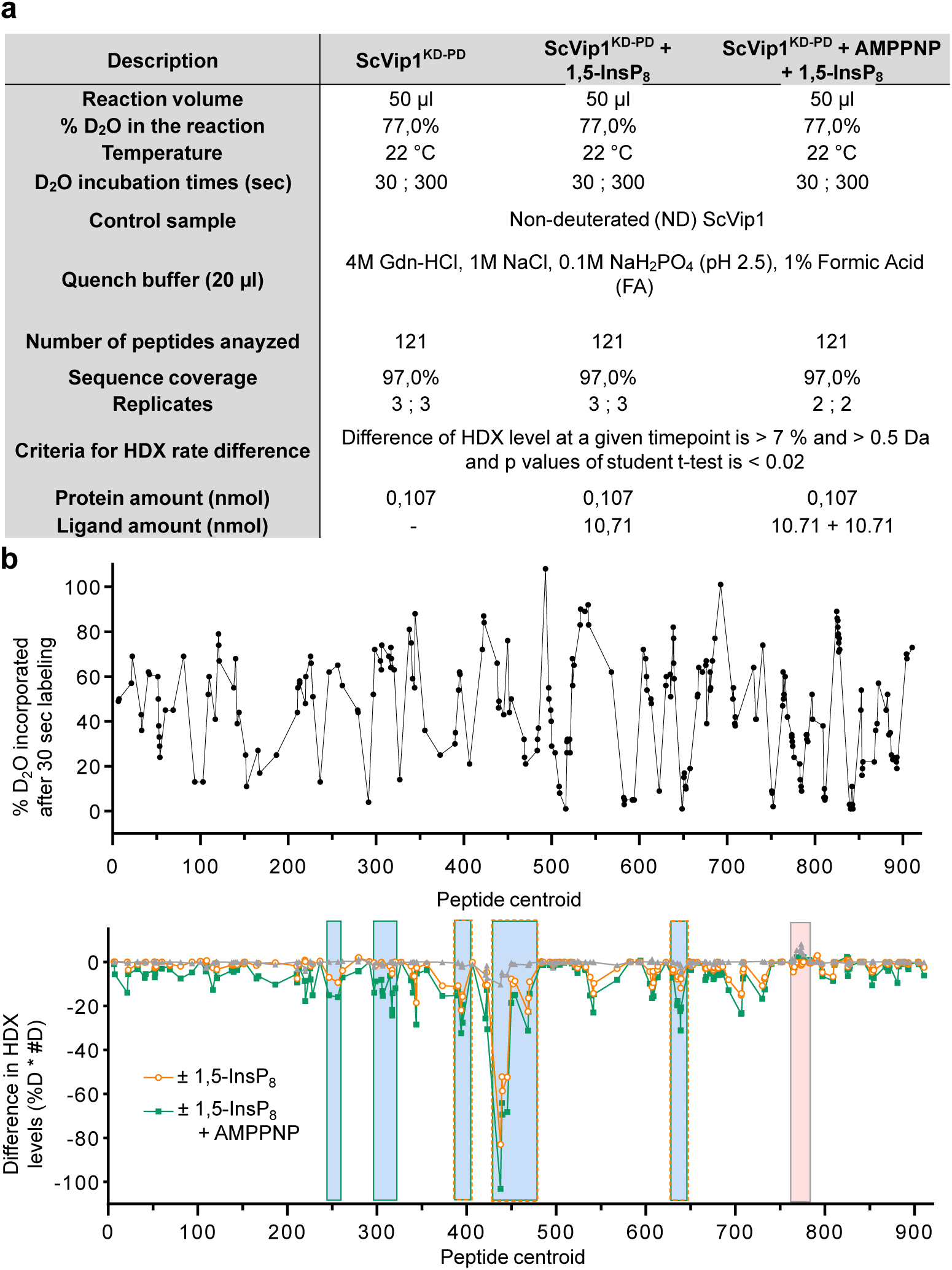
Summary of the HDX-MS experiments. **a** Experimental details of the HDX analyses. **b** D_2_O incorporation into ScVip1^KD-PD^. **c** HDX differences comparing apo ScVip1^KD-PD^, ScVip1^KD-PD^ in complex with 1,5-InsP_8_ and ScVip1^KD-PD^ in the presence of 1,5-InsP_8_ and a non-hydrolyzable ATP analog (blue, protection; red, deprotection).

**Supplementary Fig. 12.**
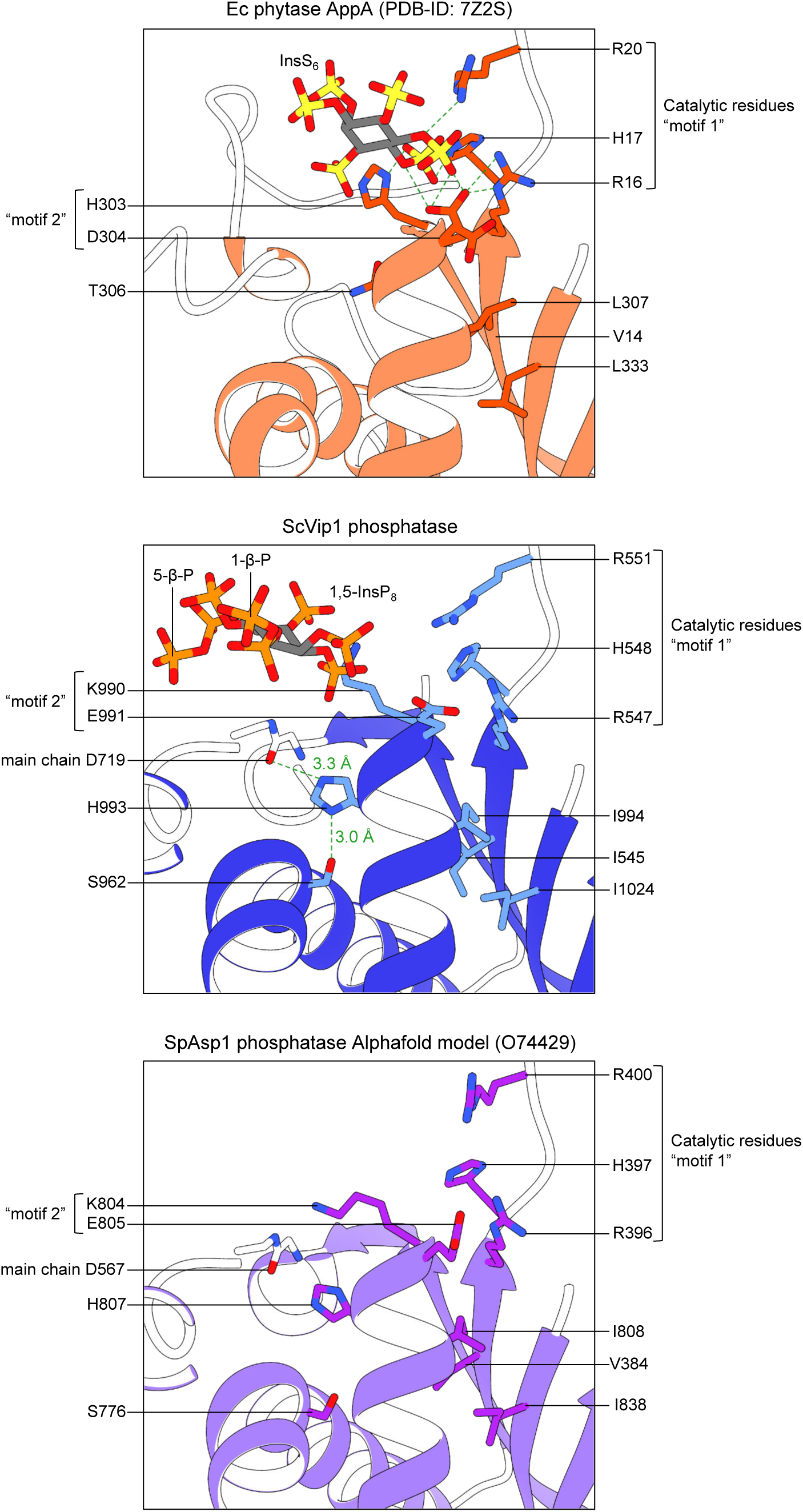
The “HD” catalytic motif presents in phytases is not present in ScVip1^PD^. Detailed views of the active sites of *E. coli* phytase AppA (in orange, PDP-ID 7Z2S^60^) bounds to the substrate analogue InsS_6_, *S. cerevisiae* Vip1^PD Δ848-918^, and the Alphafold model of *S. pombe* Asp1^PD^ (in purple, https://alphafold.ebi.ac.uk/entry/O74429)^103^. The InsP_6_ and 1,5-InsP_8_ molecules are depicted in bonds representation, and the side chains of residues belonging to the catalytic motifs 1 and 2 are depicted bonds representation. Residues His303 and Asp304 forming the HD motif in AppA correspond to Lys990 and Glu991 in ScVip1^PD^ and not to His993 and Ile994, as previously thought^23,24,35,41,42^.

**Supplementary Table 1 *Saccharomyces cerevisiae* strains and plasmids used in this study**

**Supplementary Table 2 Synthetic gene and primer sequences used in this study**

**Supplementary Table 3 Overview of transgenic Arabidopsis lines used in this study**

**Supplementary Table 4 Raw HDX-MS data**

## References

1. Shears, S. B. Intimate connections: Inositol pyrophosphates at the interface of metabolic regulation and cell signaling. Journal Cellular Physiology 233, 1897–1912 (2018).

2. Gu, C., Li, X., Zong, G., Wang, H. & Shears, S. B. IP8: A quantitatively minor inositol pyrophosphate signaling molecule that punches above its weight. Advances in Biological Regulation 91, 101002 (2024).

3. Nguyen Trung, M., Furkert, D. & Fiedler, D. Versatile signaling mechanisms of inositol pyrophosphates. Current Opinion in Chemical Biology 70, 102177 (2022).

4. Wild, R. et al. Control of eukaryotic phosphate homeostasis by inositol polyphosphate sensor domains. Science 352, 986–990 (2016).

5. Chabert, V. et al. Inositol pyrophosphate dynamics reveals control of the yeast phosphate starvation program through 1,5-IP8 and the SPX domain of Pho81. Elife 12, RP87956 (2023).

6. Guan, Z. et al. The cytoplasmic synthesis and coupled membrane translocation of eukaryotic polyphosphate by signal-activated VTC complex. Nat Commun 14, 718 (2023).

7. Cordeiro, C. D., Saiardi, A. & Docampo, R. The inositol pyrophosphate synthesis pathway in Trypanosoma brucei is linked to polyphosphate synthesis in acidocalcisomes. Mol Microbiol 106, 319–333 (2017).

8. Couso, I. et al. Synergism between Inositol Polyphosphates and TOR Kinase Signaling in Nutrient Sensing, Growth Control, and Lipid Metabolism in Chlamydomonas. Plant Cell 28, 2026–2042 (2016).

9. Stevenson-Paulik, J., Bastidas, R. J., Chiou, S.-T., Frye, R. A. & York, J. D. Generation of phytate-free seeds in *Arabidopsis* through disruption of inositol polyphosphate kinases. Proc. Natl. Acad. Sci. U.S.A. 102, 12612–12617 (2005).

10. Zhu, J. et al. Two bifunctional inositol pyrophosphate kinases/phosphatases control plant phosphate homeostasis. eLife 8, e43582 (2019).

11. Dong, J. et al. Inositol Pyrophosphate InsP8 Acts as an Intracellular Phosphate Signal in Arabidopsis. Molecular Plant (2019) doi:10.1016/j.molp.2019.08.002.

12. Ried, M. K. et al. Inositol pyrophosphates promote the interaction of SPX domains with the coiled-coil motif of PHR transcription factors to regulate plant phosphate homeostasis. Nature Communications 12, 384 (2021).

13. Riemer, E. et al. ITPK1 is an InsP6/ADP phosphotransferase that controls phosphate signaling in Arabidopsis. Molecular Plant 14, 1864–1880 (2021).

14. Zhou, J. et al. Mechanism of phosphate sensing and signaling revealed by rice SPX1-PHR2 complex structure. Nat Commun 12, 7040 (2021).

15. Guan, Z. et al. Mechanistic insights into the regulation of plant phosphate homeostasis by the rice SPX2 - PHR2 complex. Nat Commun 13, 1581 (2022).

16. Gu, C. et al. The Significance of the Bifunctional Kinase/Phosphatase Activities of Diphosphoinositol Pentakisphosphate Kinases (PPIP5Ks) for Coupling Inositol Pyrophosphate Cell Signaling to Cellular Phosphate Homeostasis. J. Biol. Chem. 292, 4544–4555 (2017).

17. Li, X. et al. Control of XPR1-dependent cellular phosphate efflux by InsP8 is an exemplar for functionally-exclusive inositol pyrophosphate signaling. PNAS 117, 3568–3574 (2020).

18. Wang, Z. et al. Rapid stimulation of cellular Pi uptake by the inositol pyrophosphate InsP _8_ induced by its photothermal release from lipid nanocarriers using a near infra-red light-emitting diode. Chem. Sci. 11, 10265–10278 (2020).

19. Li, X. et al. Homeostatic coordination of cellular phosphate uptake and efflux requires an organelle-based receptor for the inositol pyrophosphate IP8. Cell Rep 43, 114316 (2024).

20. Saiardi, A., Erdjument-Bromage, H., Snowman, A. M., Tempst, P. & Snyder, S. H. Synthesis of diphosphoinositol pentakisphosphate by a newly identified family of higher inositol polyphosphate kinases. Current Biology 9, 1323–1326 (1999).

21. Dubois, E. et al. In Saccharomyces cerevisiae, the Inositol Polyphosphate Kinase Activity of Kcs1p Is Required for Resistance to Salt Stress, Cell Wall Integrity, and Vacuolar Morphogenesis. Journal of Biological Chemistry 277, 23755–23763 (2002).

22. Laha, D. et al. Arabidopsis ITPK1 and ITPK2 Have an Evolutionarily Conserved Phytic Acid Kinase Activity. ACS Chem. Biol. 14, 2127–2133 (2019).

23. Fridy, P. C., Otto, J. C., Dollins, D. E. & York, J. D. Cloning and characterization of two human VIP1-like inositol hexakisphosphate and diphosphoinositol pentakisphosphate kinases. J. Biol. Chem. 282, 30754–30762 (2007).

24. Mulugu, S. et al. A Conserved Family of Enzymes That Phosphorylate Inositol Hexakisphosphate. Science 316, 106–109 (2007).

25. Dollins, D. E. et al. Vip1 is a kinase and pyrophosphatase switch that regulates inositol diphosphate signaling. Proceedings of the National Academy of Sciences 117, 9356–9364 (2020).

26. Gokhale, N. A., Zaremba, A. & Shears, S. B. Receptor-dependent compartmentalization of PPIP5K1, a kinase with a cryptic polyphosphoinositide binding domain. Biochemical Journal 434, 415–426 (2011).

27. Wang, H., Falck, J. R., Hall, T. M. T. & Shears, S. B. Structural basis for an inositol pyrophosphate kinase surmounting phosphate crowding. Nat Chem Biol 8, 111–116 (2012).

28. Weaver, J. D., Wang, H. & Shears, S. B. The kinetic properties of a human PPIP5K reveal that its kinase activities are protected against the consequences of a deteriorating cellular bioenergetic environment. Bioscience Reports 33, e00022 (2013).

29. Nair, V. S. et al. Inositol pyrophosphate synthesis by diphosphoinositol pentakisphosphate kinase-1 is regulated by phosphatidylinositol(4,5)bisphosphate. Bioscience Reports 38, BSR20171549 (2018).

30. Desfougères, Y. et al. The inositol pyrophosphate metabolism of Dictyostelium discoideum does not regulate inorganic polyphosphate (polyP) synthesis. Advances in Biological Regulation 83, 100835 (2022).

31. Desai, M. et al. Two inositol hexakisphosphate kinases drive inositol pyrophosphate synthesis in plants. Plant J. 80, 642–653 (2014).

32. Laha, D. et al. VIH2 Regulates the Synthesis of Inositol Pyrophosphate InsP8 and Jasmonate-Dependent Defenses in Arabidopsis. Plant Cell 27, 1082–1097 (2015).

33. Shukla, A. et al. Wheat inositol pyrophosphate kinase TaVIH2-3B modulates cell-wall composition and drought tolerance in Arabidopsis. BMC Biol 19, 261 (2021).

34. Laurent, F. et al. Inositol pyrophosphate catabolism by three families of phosphatases controls plant growth and development. 2024.04.30.591831 Preprint at 10.1101/2024.04.30.591831 (2024).

35. Pascual-Ortiz, M. et al. Asp1 Bifunctional Activity Modulates Spindle Function via Controlling Cellular Inositol Pyrophosphate Levels in Schizosaccharomyces pombe. Mol. Cell. Biol. 38, pii: e00047-18 (2018).

36. Gu, C. et al. KO of 5-InsP7 kinase activity transforms the HCT116 colon cancer cell line into a hypermetabolic, growth-inhibited phenotype. Proceedings of the National Academy of Sciences 114, 11968–11973 (2017).

37. Yousaf, R. et al. Mutations in Diphosphoinositol-Pentakisphosphate Kinase PPIP5K2 are associated with hearing loss in human and mouse. PLoS Genet 14, e1007297 (2018).

38. Gu, C. et al. Metabolic supervision by PPIP5K, an inositol pyrophosphate kinase/phosphatase, controls proliferation of the HCT116 tumor cell line. Proc. Natl. Acad. Sci. U.S.A. 118, e2020187118 (2021).

39. Gulabani, H. et al. Arabidopsis inositol polyphosphate kinases IPK1 and ITPK1 modulate crosstalk between SA-dependent immunity and phosphate-starvation responses. Plant Cell Rep 41, 347–363 (2022).

40. Benjamin, B. et al. Activities and Structure-Function Analysis of Fission Yeast Inositol Pyrophosphate (IPP) Kinase-Pyrophosphatase Asp1 and Its Impact on Regulation of *pho1* Gene Expression. mBio 13, e01034–22 (2022).

41. Pöhlmann, J. et al. The Vip1 inositol polyphosphate kinase family regulates polarized growth and modulates the microtubule cytoskeleton in fungi. PLoS Genet. 10, e1004586 (2014).

42. Wang, H. et al. Asp1 from *Schizosaccharomyces pombe* Binds a [2Fe-2S] ^2+^ Cluster Which Inhibits Inositol Pyrophosphate 1-Phosphatase Activity. Biochemistry 54, 6462–6474 (2015).

43. Gu, C., Wilson, M. S. C., Jessen, H. J., Saiardi, A. & Shears, S. B. Inositol Pyrophosphate Profiling of Two HCT116 Cell Lines Uncovers Variation in InsP8 Levels. PLOS ONE 11, e0165286 (2016).

44. Qiu, D. et al. Analysis of inositol phosphate metabolism by capillary electrophoresis electrospray ionization mass spectrometry. Nat Commun 11, 6035 (2020).

45. Wang, H. et al. Synthetic Inositol Phosphate Analogs Reveal that PPIP5K2 Has a Surface-Mounted Substrate Capture Site that Is a Target for Drug Discovery. Chemistry & Biology 21, 689–699 (2014).

46. Benjamin, B. et al. Structures of Fission Yeast Inositol Pyrophosphate Kinase Asp1 in Ligand-Free, Substrate-Bound, and Product-Bound States. mBio 13, e03087–22 (2022).

47. Rigden, D. J. The histidine phosphatase superfamily: structure and function. Biochemical Journal 409, 333–348 (2008).

48. Scheffzek, K. & Welti, S. Pleckstrin homology (PH) like domains – versatile modules in protein–protein interaction platforms. FEBS Letters 586, 2662–2673 (2012).

49. Holm, L. & Rosenström, P. Dali server: conservation mapping in 3D. Nucl. Acids Res. 38, W545–W549 (2010).

50. Aravind, L. & Ponting, C. P. The GAF domain: an evolutionary link between diverse phototransducing proteins. Trends Biochem Sci 22, 458–459 (1997).

51. Ponting, C. P. & Aravind, L. PAS: a multifunctional domain family comes to light. Curr Biol 7, R674–677 (1997).

52. Ho, Y. J., Burden, L. M. & Hurley, J. H. Structure of the GAF domain, a ubiquitous signaling motif and a new class of cyclic GMP receptor. The EMBO Journal 19, 5288–5299 (2000).

53. Henry, J. T. & Crosson, S. Ligand-Binding PAS Domains in a Genomic, Cellular, and Structural Context. Annual Review of Microbiology 65, 261–286 (2011).

54. Milburn, C. C. et al. Binding of phosphatidylinositol 3,4,5-trisphosphate to the pleckstrin homology domain of protein kinase B induces a conformational change. Biochemical Journal 375, 531–538 (2003).

55. Wang, Q. et al. Autoinhibition of Bruton’s tyrosine kinase (Btk) and activation by soluble inositol hexakisphosphate. Elife 4, e06074 (2015).

56. Lim, D., Golovan, S., Forsberg, C. W. & Jia, Z. Crystal structures of Escherichia coli phytase and its complex with phytate. Nat Struct Biol 7, 108–113 (2000).

57. Ostanin, K. et al. Overexpression, site-directed mutagenesis, and mechanism of Escherichia coli acid phosphatase. J Biol Chem 267, 22830–22836 (1992).

58. Ostanin, K. & Van Etten, R. L. Asp304 of Escherichia coli acid phosphatase is involved in leaving group protonation. J Biol Chem 268, 20778–20784 (1993).

59. Auesukaree, C. et al. Intracellular phosphate serves as a signal for the regulation of the PHO pathway in Saccharomyces cerevisiae. J. Biol. Chem. 279, 17289–17294 (2004).

60. Acquistapace, I. M. et al. Insights to the Structural Basis for the Stereospecificity of the Escherichia coli Phytase, AppA. IJMS 23, 6346 (2022).

61. Stentz, R. et al. A Bacterial Homolog of a Eukaryotic Inositol Phosphate Signaling Enzyme Mediates Cross-kingdom Dialog in the Mammalian Gut. Cell Reports 6, 646–656 (2014).

62. Pan, X. et al. A robust toolkit for functional profiling of the yeast genome. Mol Cell 16, 487–496 (2004).

63. Masinas, M. P. D., Usaj, M. M., Usaj, M., Boone, C. & Andrews, B. J. TheCellVision.org: A Database for Visualizing and Mining High-Content Cell Imaging Projects. G3 (Bethesda) 10, 3969–3976 (2020).

64. Schwartz, K., Richards, K. & Botstein, D. BIM1 encodes a microtubule-binding protein in yeast. Mol Biol Cell 8, 2677–2691 (1997).

65. Manatschal, C. et al. Molecular basis of Kar9-Bim1 complex function during mating and spindle positioning. Mol Biol Cell 27, 3729–3745 (2016).

66. An, Y., Jessen, H. J., Wang, H., Shears, S. B. & Kireev, D. Dynamics of Substrate Processing by PPIP5K2, a Versatile Catalytic Machine. Structure 27, 1022–1028.e2 (2019).

67. Sanchez, A. M., Schwer, B., Jork, N., Jessen, H. J. & Shuman, S. Activities, substrate specificity, and genetic interactions of fission yeast Siw14, a cysteinyl-phosphatase-type inositol pyrophosphatase. mBio 14, e02056–23 (2023).

68. Ghosh, S. et al. Activities and genetic interactions of fission yeast Aps1, a Nudix-type inositol pyrophosphatase and inorganic polyphosphatase. mBio 15, e0108424 (2024).

69. Garg, A., Shuman, S. & Schwer, B. A genetic screen for suppressors of hyper-repression of the fission yeast PHO regulon by Pol2 CTD mutation T4A implicates inositol 1-pyrophosphates as agonists of precocious lncRNA transcription termination. Nucleic Acids Res 48, 10739–10752 (2020).

70. Wang, H., Gu, C., Rolfes, R. J., Jessen, H. J. & Shears, S. B. Structural and biochemical characterization of Siw14: A protein-tyrosine phosphatase fold that metabolizes inositol pyrophosphates. Journal of Biological Chemistry 293, 6905–6914 (2018).

71. Lee, Y.-H., Li, Y., Uyeda, K. & Hasemann, C. A. Tissue-specific Structure/Function Differentiation of the Liver Isoform of 6-Phosphofructo-2-kinase/Fructose-2,6-bisphosphatase. Journal of Biological Chemistry 278, 523–530 (2003).

72. Machkalyan, G., Trieu, P., Pétrin, D., Hébert, T. E. & Miller, G. J. PPIP5K1 interacts with the exocyst complex through a C-terminal intrinsically disordered domain and regulates cell motility. Cellular Signalling 28, 401–411 (2016).

73. Hothorn, M., Dabi, T. & Chory, J. Structural basis for cytokinin recognition by Arabidopsis thaliana histidine kinase 4. Nat. Chem. Biol. 7, 766–768 (2011).

74. Hashimoto, Y., Zhang, S. & Blissard, G. W. Ao38, a new cell line from eggs of the black witch moth, Ascalapha odorata (Lepidoptera: Noctuidae), is permissive for AcMNPV infection and produces high levels of recombinant proteins. BMC Biotechnol. 10, 50 (2010).

75. Malakhov, M. P. et al. SUMO fusions and SUMO-specific protease for efficient expression and purification of proteins. J. Struct. Funct. Genomics 5, 75–86 (2004).

76. Gorrec, F. The MORPHEUS protein crystallization screen. J Appl Crystallogr 42, 1035–1042 (2009).

77. Kabsch, W. Automatic processing of rotation diffraction data from crystals of initially unknown symmetry land cell constants. Journal of Applied Crystallography 26, 795–800 (1993).

78. McCoy, A. J. et al. Phaser crystallographic software. J. Appl. Crystallogr. 40, 658–674 (2007).

79. Emsley, P. & Cowtan, K. Coot: model-building tools for molecular graphics. Acta Cryst D 60, 2126–2132 (2004).

80. Afonine, P. V. et al. Towards automated crystallographic structure refinement with phenix.refine. Acta Cryst D 68, 352–367 (2012).

81. Read, R. J. & McCoy, A. J. Using SAD data in Phaser. Acta Crystallogr D Biol Crystallogr 67, 338–344 (2011).

82. Terwilliger, T. C. SOLVE and RESOLVE: automated structure solution and density modification. Meth. Enzymol. 374, 22–37 (2003).

83. Lebedev, A. A. & Isupov, M. N. Space-group and origin ambiguity in macromolecular structures with pseudo-symmetry and its treatment with the program Zanuda. Acta Cryst D 70, 2430–2443 (2014).

84. Ban, N. et al. Placement of protein and RNA structures into a 5 A-resolution map of the 50S ribosomal subunit. Nature 400, 841–847 (1999).

85. Wittmann, J. G. & Rudolph, M. G. Pseudo-merohedral twinning in monoclinic crystals of human orotidine-5’-monophosphate decarboxylase. Acta Crystallogr D Biol Crystallogr 63, 744–749 (2007).

86. Davis, I. W. et al. MolProbity: all-atom contacts and structure validation for proteins and nucleic acids. Nucleic Acids Res 35, W375–383 (2007).

87. Pettersen, E. F. et al. UCSF Chimera--a visualization system for exploratory research and analysis. J Comput Chem 25, 1605–1612 (2004).

88. Harmel, R. K. et al. Harnessing 13C-labeled myo-inositol to interrogate inositol phosphate messengers by NMR. Chem Sci 10, 5267–5274 (2019).

89. Ames, B. Assay of inorganic phosphate, total phosphate and phosphatases. Methods Enzymol 8, 115–118 (1966).

90. Baykov, A. A., Evtushenko, O. A. & Avaeva, S. M. A malachite green procedure for orthophosphate determination and its use in alkaline phosphatase-based enzyme immunoassay. Anal. Biochem. 171, 266–270 (1988).

91. Longtine, M. S. et al. Additional modules for versatile and economical PCR-based gene deletion and modification in Saccharomyces cerevisiae. Yeast 14, 953–961 (1998).

92. Binder, A. et al. A modular plasmid assembly kit for multigene expression, gene silencing and silencing rescue in plants. PLoS One 9, e88218 (2014).

93. Clough, S. J. & Bent, A. F. Floral dip: a simplified method for Agrobacterium-mediated transformation of Arabidopsis thaliana. The Plant Journal 16, 735–743 (1998).

94. Schindelin, J. et al. Fiji: an open-source platform for biological-image analysis. Nat. Methods 9, 676–682 (2012).

95. Punjani, A., Rubinstein, J. L., Fleet, D. J. & Brubaker, M. A. cryoSPARC: algorithms for rapid unsupervised cryo-EM structure determination. Nat Methods 14, 290–296 (2017).

96. Wang, H. et al. Autoregulation of Class II Alpha PI3K Activity by Its Lipid-Binding PX-C2 Domain Module. Mol Cell 71, 343–351.e4 (2018).

97. Hostachy, S. et al. Dissecting the activation of insulin degrading enzyme by inositol pyrophosphates and their bisphosphonate analogs. Chem Sci 12, 10696–10702 (2021).

98. Perez-Riverol, Y. et al. The PRIDE database resources in 2022: a hub for mass spectrometry-based proteomics evidences. Nucleic Acids Res 50, D543–D552 (2022).

99. Ortlund, E., LaCount, M. W. & Lebioda, L. Crystal Structures of Human Prostatic Acid Phosphatase in Complex with a Phosphate Ion and α-Benzylaminobenzylphosphonic Acid Update the Mechanistic Picture and Offer New Insights into Inhibitor Design,. Biochemistry 42, 383–389 (2003).

100. Hamada, K. et al. Crystal structure of the protein histidine phosphatase SixA in the multistep His-Asp phosphorelay. Genes to Cells 10, 1–11 (2005).

101. Landau, M. et al. ConSurf 2005: the projection of evolutionary conservation scores of residues on protein structures. Nucleic Acids Res 33, W299–302 (2005).

102. Dunnett, C. W. A Multiple Comparison Procedure for Comparing Several Treatments with a Control. Journal of the American Statistical Association 50, 1096–1121 (1955).

103. Varadi, M. et al. AlphaFold Protein Structure Database in 2024: providing structure coverage for over 214 million protein sequences. Nucleic Acids Research 52, D368–D375 (2024).

